# Granule cells reorient cortical trajectories to separate contexts

**DOI:** 10.64898/2026.03.03.709240

**Authors:** Martha G. Garcia-Garcia, Michał J. Wójcik, Srijan Thota, Luke Drake, Amma Otchere, Oluwatobi Akinwale, Lizmaylin Ramos, Rui Ponte Costa, Mark J. Wagner

## Abstract

To learn effectively, animals must generalize across yet distinguish between related contexts. Generalization relies on low-dimensional neural manifolds found throughout the neocortex^1,2^, which accelerate learning by constraining neural activity to task-relevant axes^3^. Conversely, context separation is attributed to neural expansion layers that can project information into high-dimensional feature spaces^4,5^, most famously cerebellar granule cells (GrCs)^6–8^. To investigate the generalization-separation tradeoff, we simultaneously imaged key nodes in the universal cortico-cerebellar pathway^9^—premotor layer 5 pyramidal tract (L5PT) and GrCs—during parallel learning of two distinct skills with shared temporal structure. Rather than expanding the cortical representations, GrCs retained their low-rank encoding of each task. Across contexts, despite stable cortico-cerebellar coupling, L5PT activity patterns generalized while GrC patterns temporally remapped. But rather than independently scrambling, GrC populations remapped coherently: their low-dimensional trajectories “rotated” apart between tasks, separating the contexts while preserving the cortical geometry of each. Moreover, GrC trajectories diverged most strongly in expert animals. This suggests a fundamental architectural division of labor: the cortex provides invariant dynamic primitives for smooth generalization, while cerebellar activity reconfigures them to drive context-specific output.

## Main

Generalization allows animals to accelerate new learning guided by past experience. In the brain, however, using the vast combinatorial repertoire of theoretically possible neural activity patterns would make generalization and rapid learning computationally prohibitive, creating a “curse of dimensionality”. To solve this dilemma, the neocortex employs low-dimensional neural manifolds^1,3^—population activity patterns constrained to a low-rank subspace in a high-dimensional embedding. By restricting activity to axes capturing essential behavioral variables (e.g., time, speed), manifolds help circuits generalize to previously unseen task variations^10^. They can also facilitate generalization *across* contexts through “structural learning”^11^: using the underlying features of one task to quickly learn another. These advantages strongly incentivize reusing manifolds to make circuits robust^12^ to both within- and cross-context variations. Accordingly, although the cortex can separate related contexts when needed^13^, evidence suggests that whenever possible, the brain prioritizes repurposing low-dimensional neural structures^2,3,14^—a fundamental property shared with artificial neural networks^15,16^.

However, manifold reuse comes at a steep cost: interference^17^. When two contexts share overlapping neural activity patterns, it is difficult to drive distinct downstream control policies^18^. A classical architecture for distinguishing contexts is the neural “expansion layer”: by projecting overlapping inputs into a high-rank feature space^4,5^, patterns can be maximally separated^6–8^. This motif is ubiquitous in classification circuits: overlapping odor representations are discretized into sparse clusters in piriform cortex^19^; smooth representations of visual space transform into categorical objects in inferior temporal cortex^20^; continuous encoding of physical space is converted to a contextual “hash code” in dentate gyrus^21^. Most famously, the greatest capacity for expansion is found in cerebellar granule cells (GrC), which collectively outnumber all other neurons in the brain combined^22^. Yet, while GrCs possess the capacity for high-rank representations^23^, this presents a paradox for manifolds: using high-dimensional expansion to maximally separate points along a low-dimensional trajectory precludes the smooth geometry required for continuous dynamic prediction and structural learning^24,25^.

These opposing constraints create a fundamental generalization-separation tradeoff: low-dimensional manifolds solve the “curse of dimensionality” but invite interference; high-dimensional expansion solves interference but exacerbates the “curse”. A direct interface between these competing demands is the universal cortico-cerebellar pathway that connects the layer 5 pyramidal tract (L5PT) to cerebellar GrCs via the pons^9,26^. To investigate how this circuit resolves this tradeoff, we simultaneously imaged premotor L5PTs and GrCs during “parallel skill learning,” where animals alternately trained on two tasks with distinct sensorimotor elements but shared temporal structure.

## Results

### Imaging parallel learning of two tasks

High-level task representations are shared across the rodent premotor cortex and its disynaptic GrC targets in contralateral cerebellar lobules CrusI, CrusII, and simplex^27,28^. We therefore developed simultaneous dual-site imaging of premotor L5PT and GrCs. To image L5PT neurons, we used the red-shifted Ca^2+^ indicator jRGECO1a^29^, excited with a 1064 nm laser for deeper tissue penetration (∼700 μm, **Extended Data Fig. 1a**). To specifically target L5PT, we injected AAVretro-FLP into the right pontine nuclei and AAV-fDIO-jRGECO1a into the right premotor cortex (**Fig. 1a**). We concurrently expressed the green Ca^2+^ indicator GCaMP6f in all cerebellar GrCs via the triple transgenic *Math1-Cre×Ai93×ztTA* (**Fig. 1b**). Using a custom dual-site two-photon microscope, we visualized both populations through contralateral cranial windows (**Video S1**).

**Figure 1.**
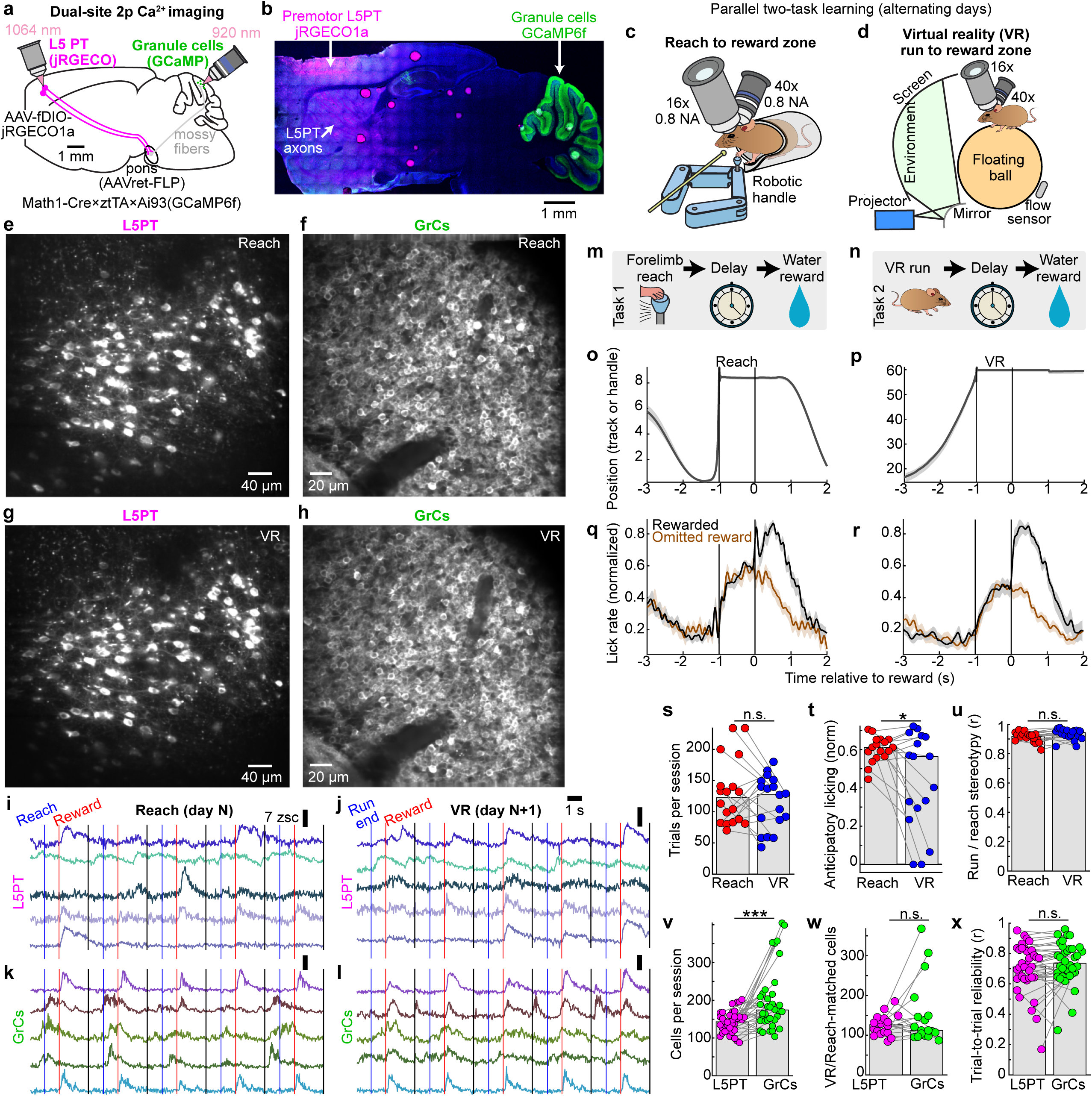
| Simultaneous cortex and cerebellar 2p imaging during parallel learning of two novel skills. **a,b,** Dual-site imaging strategy. **a**, Schematic of two-photon imaging of right premotor cortex pons-projecting layer 5 pyramidal tract neurons (L5PT) and cerebellar granule cells (GrCs) in left Crus I/II or simplex. **b**, Histology. L5PTs labeled with jRGECO1a (via AAVretro-FLP in pons and AAV-fDIO-jRGECO1a in cortex) and GrCs labeled with GCaMP6f (via Math1-Cre×Ai93×ztTA transgenics). **c,d,** The Reach-for-reward task (**c**) and virtual reality (VR) run-for-reward task (**d**), imaged with a custom dual-site microscope (**Extended Data Fig. 1**). **e–h,** Representative fields of view showing populations of L5PTs (**e,g**) and GrCs (**f,h**) tracked across consecutive Reach and VR sessions. **i-l**, Example simultaneous activity traces from five tracked L5PTs and GrCs during Reach and VR trials. Vertical lines indicate movement offset (blue), reward delivery (red), and trial end (black). **m,n,** Trial structure. Both tasks shared a timeline: action initiation (Reach vs. Run), delay, reward consumption and inter-trial interval. **o-r**, Behavioral performance. Cohort-averaged position trajectories of the reach handle (**o**) or running track position (**p**), and corresponding lick rates (**q,r**; normalized per-session, **Methods**). Shaded regions denote s.e.m. **s-u**, Task comparability. In Reach and VR, animals executed matched numbers of trials **(s),** with broadly similar levels of anticipatory licking (mean rate [-0.5, 0] s relative to reward; **t**) and movement stereotypy (**u**; p=0.3, 0.04, and 0.09). These and all subsequent bars denote medians. **v-x**, Imaging statistics. **v,** Total active cells per session (p=0.0005). **w**, Count of cells active, reliable, and tracked across both tasks (p=0.4). **x**, Response reliability (correlation of odd vs. even trial averages, per-session medians across cells; p=0.055). **o-x** n=18 matched VR and Reach sessions from 9 mice, except licking **q,t** which had only 17. All paired comparisons via Wilcoxon signed-rank test.

To investigate the neural mechanisms of task generalization versus separation, we designed a paradigm where water-restricted mice learned two tasks in parallel: a virtual reality (“VR”) run-for-reward task^30^ and a robotic manipulandum “Reach”-for-reward task^31^ (**Fig. 1c,d**), alternating between contexts over a median of 18 days (range: 11-22). The behavioral apparatuses were engineered to be physically compatible with the microscope geometry, enabling us to revisit the same imaging fields and track neural populations across behavioral contexts (**Fig. 1e-l, Extended Data Fig. 1b-h**). In the Reach task, animals stood in a tube and used their left forepaw to push a robotic manipulandum to a threshold distance of 6 mm (maximum extent: 8 mm), after which they waited through a 1-s delay to receive a water reward. Following a 2-s consumption window, the robot automatically returned to the start position. In the VR task, animals stood on an air-suspended ball and self-initiated locomotion through a virtual linear environment to a target zone 60 mm away. Upon reaching the target, the ball locked in place and animals waited through an identical 1-s delay for reward. After reward delivery, the screen darkened during a 2-s consumption and inter-trial interval (ITI) before repopulating at the start location (**Video S2**).

The sensorimotor differences between contexts were extensive: tube-constrained versus ball-restrained; dark room versus bright virtual environment; single-limb control versus whole-body locomotion; and manipulation of a handle versus a sphere. Despite these differences, the tasks shared a rigid temporal scaffold: Action→1-s Delay→Reward+ITI (**Fig. 1m,n**). In both contexts, animals learned stereotyped movement trajectories, either of the robotic handle (**Fig. 1o**) or the ball rotation (**Fig. 1p**). Furthermore, animals developed predictive licking time-locked to reward delivery, even on unexpected reward omission trials (**Fig. 1q,r**; 20% omission, randomly interleaved). In both tasks, animals executed comparable numbers of trials (**Fig. 1s**, Reach: 123±16, VR 128±17), developed comparable anticipatory licking that was slightly higher in Reach (**Fig. 1t**, Reach: 0.61±0.02, VR: 0.56±0.1, p=0.04), and achieved similar movement stereotypy (**Fig. 1u**, Reach r=0.92±0.01, VR r=0.94±0.01; data from 9 mice; these and subsequent quantifications median±s.e._median_, bootstrapped). Importantly, optogenetic inhibition confirmed that normal GrC activity was required for anticipatory licking in both tasks (**Extended Data Fig. 2**).

We reasoned that different neural circuits might use one of three strategies to encode the two behaviors:

1. Generalization, which uses overlapping neural trajectories to support the transfer of learned timing rules^2,3^.
2. Separation through high-rank expansion, which orthogonalizes neural states^32^ within and across contexts but sacrifices smoothness by distorting local Euclidean neighborhood relationships^33–35^.
3. Structured separation, which preserves representational geometry but reorients trajectories apart within the shared neural state space.

### Tracking L5PTs and GrCs across tasks

To compare L5PT and GrC dynamics during the learning of two tasks, we tracked individual neurons across matched session pairs, consisting of either cross-task transitions (18 matched VR and Reach session-pairs, typically from consecutive days) or, as a control, same-task transitions (9 matched VR-VR or Reach-Reach session-pairs, typically separated by 2 days; **Methods**). We applied a strict two-step filter to isolate genuine biological remapping from technical artifacts. First, to avoid conflating biological silence with technical dropout, we restricted our analysis to cells with detected activity on both days of each pair. Cross-task retention rates were inherently lower for the densely packed, ∼5-µm GrCs (58±5%) than for the sparser, ∼25-µm L5PTs (80±2%). Crucially, however, these matched the baseline rates in same-task control pairs (56±8% GrCs, 84±4% L5PT), indicating that tracking efficiency was a behavior-independent noise floor (**Extended Data Fig. 1c**).

Second, to filter out noisy or task-unrelated cells, we calculated a “reliability index” for each cell by correlating its trial-averaged activity on odd versus even trials. We included only cells that were robustly task-locked (*r* >0.4) on *both* days. This criterion demonstrated that most cells participated reliably in both contexts (**Extended Data Fig. 1d;** L5PT: 68±5%; GrCs: 77±7%). By restricting our analysis to “jointly reliable” populations, we established a rigorous baseline: if a cell time locks in both tasks *and* changes its firing pattern, it represents a genuine contextual remap. Overall, this yielded balanced datasets permitting fair comparisons between L5PTs and GrCs. We recorded comparable total cell numbers per session (**Fig. 1v**, 143±6 L5PT, 175±10 GrCs; *n=*36 sessions), registered similar numbers of active cells across tasks (**Fig. 1w**, 127±9 L5PT, 112±13 GrCs; *n=*18 session pairs), and observed equivalent reliability in both populations (**Fig. 1x**, *r_odd:even_*=0.71±0.03 L5PT, 0.74±0.03 GrCs; *n=*36 total sessions).

### Similar L5PT and GrC representations

We first compared L5PT to GrCs in their representations of each task in dual-trained mice. Trial-averaged activity profiles in the VR and Reach tasks appeared grossly similar for both cell types (**Fig. 2a-d**). We quantified this similarity using two metrics. First, the distribution of peak activity times was highly conserved between L5PT and GrCs (**Fig. 2e,f**). In both populations and contexts, ∼25% of neurons peaked during the delay period, and ∼50% peaked after reward delivery. Furthermore, L5PT and GrCs shared context-specific features, such as the ramping distribution of peak times over the prolonged VR running period, contrasted with a flatter distribution prior to the brief, ballistic Reach. Second, we computed the width at half-max of the trial-averaged activity peaks (**Fig. 2g**). Distributions were similar across cell types; small differences in duration (which reached significance in Reach) were likely because jRGECO1a has slower decay kinetics than GCaMP6f (530 ms and 140 ms^29,36^). Thus, L5PT and GrC activity was comparably dense spatially and temporally^37,38^.

**Figure 2.**
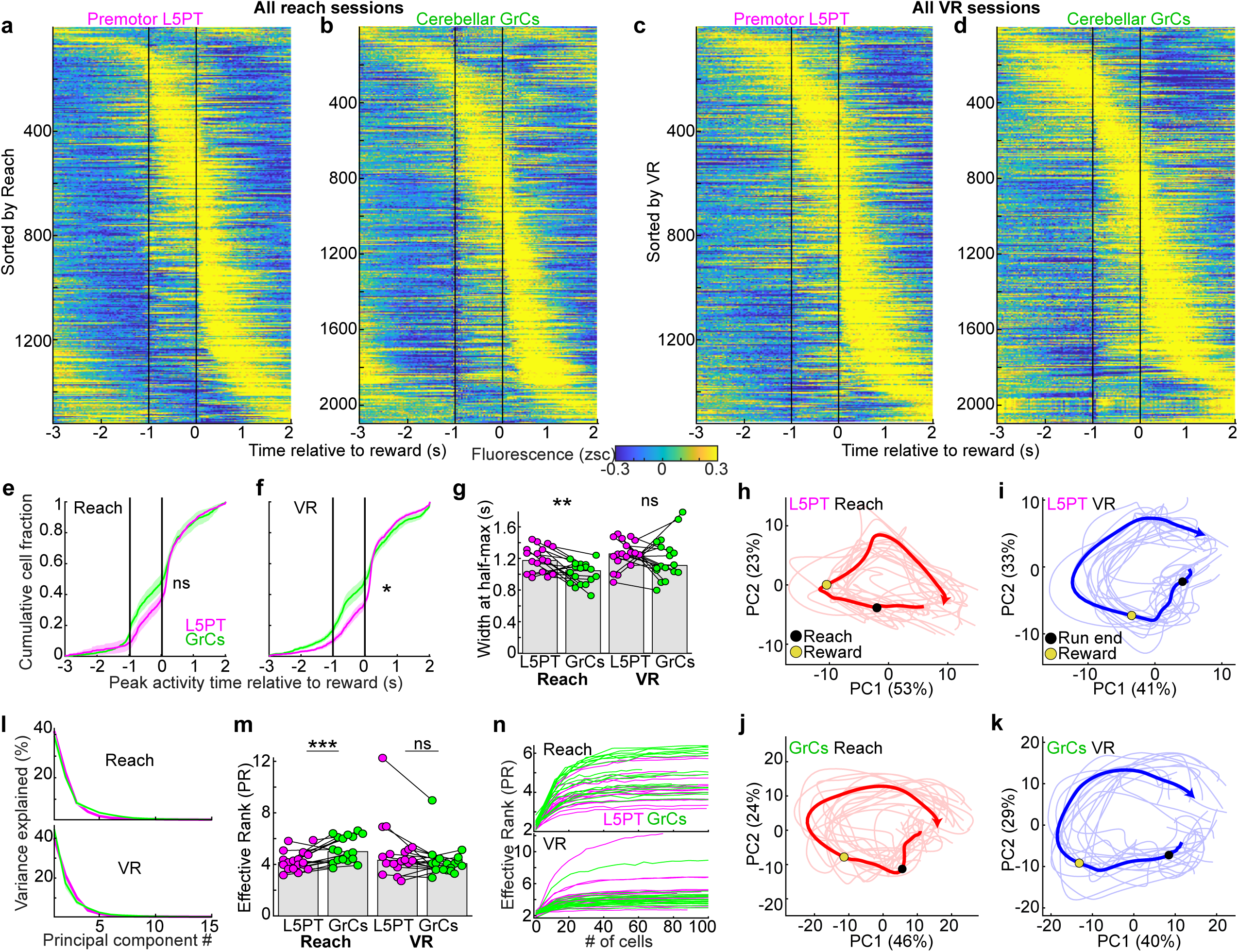
| VR and Reach tasks recruit common neural populations into similar low-dimensional dynamics. **a-d**, Task-aligned activity rasters, including all neurons active and reliable in both tasks (from 18 matched Reach-VR session pairs in 9 trained mice). Rows show fluorescence averaged across rewarded trials aligned to reward delivery, with Reach/run offset at −1 s. Cells are sorted by peak activity time independently for each panel (1,521 L5PT, 2,096 GrCs). **e,f,** Cumulative distributions of peak activity times relative to reward (mean±s.e.m. across sessions). L5PT and GrCs were indistinguishable in Reach (p=0.12), while VR featured slightly more pre-reward GrC peaks (p=0.02; permutation test, **Methods**). **g,** Temporal widths of activity peaks were slightly narrower for GrCs than L5PT in Reach but indistinguishable in VR (dots show the median full width at half maximum of trial-averaged activity for each session. Reach, p=0.006, VR, p=0.4, Wilcoxon signed-rank test). **h-k**, Representative low-dimensional population dynamics. 2D projections of neural activity onto the first 2 PCs (computed independently for each cell type and task in one Reach-VR session pair). Thick lines show the mean and thin lines show 15 single trials with the lowest deviation from the mean. Axis percentages display variance explained. Arrowheads indicate the end of the plotted temporal epoch. **l-n**, Dimensionality analysis. **l**, Scree plots show variance explained by each PC (mean±s.e.m. across 18 sessions each). **m**, Effective rank (participation ratio: (∑ *λ_i_*)^2^⁄∑ *λ_i_*^2^) for each session. GrC rank was slightly higher in Reach (p=0.001) but trended lower than L5PT in VR (p=0.054, Wilcoxon signed-rank test). **n,** Rank saturation (computed on populations subsampled at 20 intervals and averaged across 50 bootstraps). Curves saturated at cell counts far below total population size, indicating underlying low-dimensional structure. All quantifications across 18 session pairs from 9 mice.

We next considered the population structure of neural activity. Due to their high embedding dimensionality (i.e., immense population size), GrCs are postulated to separate input patterns through high-rank expansion^5–8,32^. To test whether GrC activity actually reflects such an expansion of cortical dynamics, we performed trial-averaged principal components analysis (PCA) on each task independently. Visualizing the trajectories of the top 2 principal components (PCs) revealed that both L5PT and GrC activity typically evolved along similarly cyclic, low-dimensional trajectories in both tasks (**Fig. 2h-k**). To quantify this similarity, we computed the variance explained by each PC. The scree plots were broadly similar for L5PT and GrCs, consistent with comparably low-rank dynamics (**Fig. 2l**). To estimate the effective dimensionality (i.e., rank), we computed the “participation ratio” (PR). L5PT and GrC activity exhibited similarly low effective rank, with GrCs slightly higher in Reach and trending lower in VR (**Fig. 2m**, Reach: L5 4.1±0.2, GrCs 5.0±0.5, p=0.001; VR: L5PT 4.3±0.3, GrCs 4.1±0.2, p=0.054). Computed on single-trial data, PR was significantly lower for GrCs than L5PT in both tasks (**Extended Data Fig. 3a**). To confirm that this shared low apparent rank was not an artifact of subsampling, we computed PR saturation curves and found that estimates saturated at cell counts well below the total population size (**Fig. 2n**). This conservation of rank is qualitatively inconsistent with a traditional high-dimensional orthogonalizing expansion^32,39^.

Critically, low effective rank implies a substantial compression of the available sensory and motor state space. In VR, for example, while the animal’s behavior involves high-dimensional degrees of freedom—including whole-body coordination for balance and locomotion, whisker kinematics, orofacial movements, and dense optic flow—the majority of neural variance is concentrated across a far smaller set of axes. This suggests that despite the anatomical opportunity to expand these high-dimensional sensorimotor features, GrCs instead preserved the low-rank cortical structure, failing to amplify the vast available degrees of freedom. Overall, two tasks learned in parallel with common temporal structure but widely differing sensorimotor contexts elicited population neural representations with grossly similar low-rank structure in both L5PT and GrCs.

### L5PTs generalize but GrCs remap

We next compared the responses of individual neurons across tasks. Surprisingly, individual L5PT neurons exhibited strong cross-context generalization, regardless of their peak response times (**Fig. 3a-c**). In contrast, GrCs displayed robust cross-context temporal remapping. Individual GrCs exhibited complex and heterogeneous forms of remapping, including bidirectional phase shifts and polarity inversions (**Fig. 3d-f**). Consequently, when neurons were sorted by their activity in one task, L5PTs largely maintained their sequential ordering in the opposing task, whereas GrC sequences appeared scrambled (**Fig. 3g-n**). To quantify generalization, we computed the correlation between each cell’s trial-averaged activity in VR and Reach, correcting for signal attenuation (**Methods**, reliability distributions were equivalent between cell types, **Fig. 1x**). Across recordings, L5PT cross-task activity correlations more than doubled those of GrCs (**Fig. 3o**, r=0.58±0.05 vs 0.23±0.1). Crucially, the cell types were indistinguishable in “same-task” cross-day comparisons (**Fig. 3p**). GrCs were thus more likely to remap activity across contexts, despite remaining active and reliable in both.

**Figure 3.**
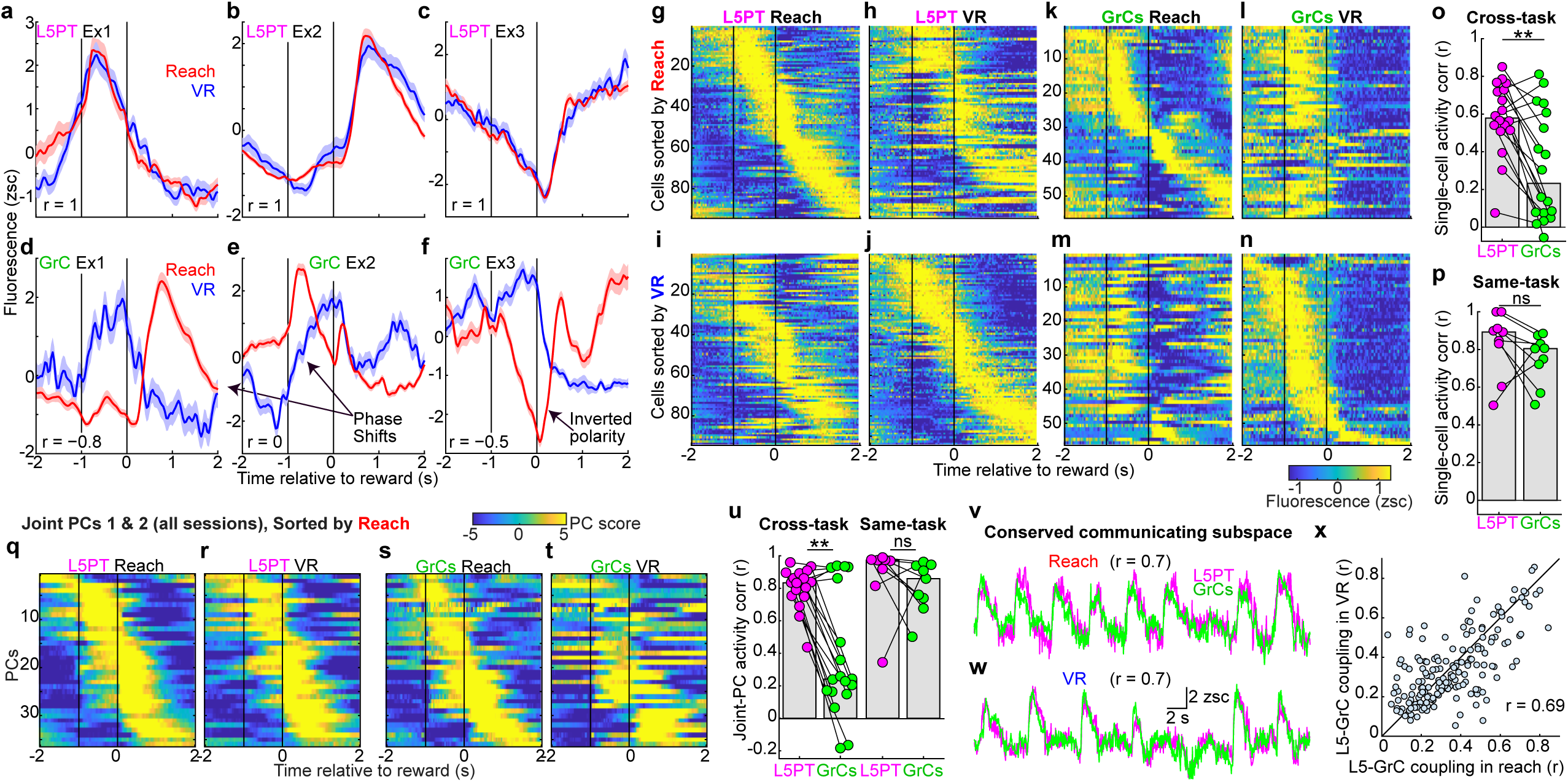
| L5PTs generalize across contexts but GrCs temporally remap. **a-f,** Representative single neurons tracked across tasks from one session pair. Traces and shaded regions show fluorescence mean±s.e.m. across rewarded trials (186 Reach and 120 VR trials). **a-c,** L5PTs exhibiting delay-ramping (**a**), reward-activated (**b**), and delay-suppressed (**c**) dynamics that generalized across contexts. **d-f,** Simultaneously-recorded GrCs that temporally remapped by phase-shifting (**d,e**) or inverting polarity (**f**). In these and all subsequent analyses, each cell’s trial-averaged activity was z-scored. For visualization only, traces were Gaussian smoothed with *σ*=40 ms. **g-n**, Population activity rasters for all cells active and reliable in both contexts from the session-pair in **a-f** (94 L5PTs and 56 GrCs). Cells are sorted by peak activity time in Reach or VR. The L5PT population temporal sequence is similar across contexts, whereas GrC sequences appear scrambled. **o,p,** Quantification of cross-task neural correlations. Pearson correlation of trial-averaged activity profiles for each cell across days (dots show session medians). L5PTs showed higher cross-task similarity than GrCs (p=0.002), but were indistinguishable in same-task control comparisons (p=0.3, Wilcoxon signed-rank test). **q-t**, Joint-task population dynamics. Rasters show the top 2 joint-task PCs (18 session pairs). Sorting by activity peak time in Reach revealed that L5PT PCs generalized across contexts while GrC PCs temporally remapped. **u,** Cross-day correlation of the top 2 joint-task PC activity profiles. L5PT population dynamics were substantially more correlated across tasks than GrCs (p=0.002), while same-task controls showed no difference (p=0.3, Wilcoxon signed-rank test). Dots show average correlations across PCs 1 and 2 per session pair. **v-x,** L5-GrC communicating subspaces (canonical correlations analysis [CCA]). **v,w,** Representative time-series of the first canonical variable (CV1) of a VR-Reach session pair. **x,** Correlation strength of CVs was conserved across Reach vs VR (173 CVs with r_L5-GrC_>0.1; *r* = 0.69, p<10^-6^, Pearson correlation). All cohort quantifications across 18 cross-task (9 mice) and 9 same-task (7 mice) session pairs.

We next considered how single-cell differences manifested at the population level. To identify neural ensembles with coherent activations across contexts, we performed ‘joint-task’ dimensionality reduction. We concatenated both tasks’ trial-averaged activity into a single joint matrix and performed PCA^40^. This approach isolates components that capture variance in both tasks, even if their temporal profiles remap. Importantly, the top two joint components captured the vast majority of variance explained by per-task PCA (L5 ∼85%, GrCs ∼80%) and exhibited equivalent cross-task stability for both cell types (**Extended Data Fig. 3b-d**). Despite this similar projection quality, visualizing the time-courses of the top joint components revealed a stark contrast: while the activity of L5PT modes appeared nearly identical across tasks, GrC modes robustly remapped (**Fig. 3q-t**, PCs 1-2 from all session pairs). Quantitatively, PC cross-task correlations in L5PT more than tripled those for GrCs (**Fig. 3u**, r=0.83±0.02 vs 0.24±0.1; indistinguishable in same-task controls).

This highlighted another fundamental distinction: for L5PT, extracting population modes substantially *increased* cross-task consistency compared to single neurons (rising from *r*=0.58 to 0.83, **Fig. 3o** vs **3u** “cross-task”, p=0.0005), implying that cross-context differences were orthogonal to the primary population axes. In sharp contrast, the same population-level analysis failed to rescue cross-context correlations in GrCs (*r*=0.23 vs 0.24, p=0.7). This demonstrates that the neural ensembles composing the dominant GrC modes collectively remapped. Finally, we tested if GrC remapping stemmed from unstable coupling to the cortex. Using joint-task canonical correlations analysis (CCA), we found that “communicating subspaces” between L5PT and GrCs remained comparably correlated across both tasks (**Fig. 3v-x**). This confirms that, despite a stable, high-fidelity core of cortico-cerebellar coupling, cross-task representations were decorrelated for GrCs, diverging sharply from their overlapping L5PT counterparts.

### GrC trajectories reorient between tasks

Together, these results raised an apparent contradiction: if each GrC’s activity remapped independently of the rest of the network (as suggested by the “scrambled” heatmaps, **Fig. 3l,m**), the resulting VR and Reach population trajectories would occupy largely distinct subspaces. Instead, the high variance explained by the joint PCs (**Extended Data Fig. 3b**) implied that the cross-context GrC remapping was coherent across neurons. Thus, to determine the population-level geometry of the remapping, we compared 2D trajectories in *independent task spaces* (revealing each task’s intrinsic low-dimensional geometry) versus the joint-task “*shared space*” (revealing their relative geometric orientation).

In independent task spaces, GrC trajectories preserved specific geometric features present in L5PT. For instance, in “Example 1,” both populations formed a ring with a movement “concavity” in Reach, and a “triangular” cycle in VR (**Fig. 4a-d**). Similarly, in “Example 2” VR, both populations exhibited an outward spiral motif (**Fig. 4g-j**). Thus, within each task, GrCs matched the cortex with cyclic low-dimensional trajectories.

**Figure 4.**
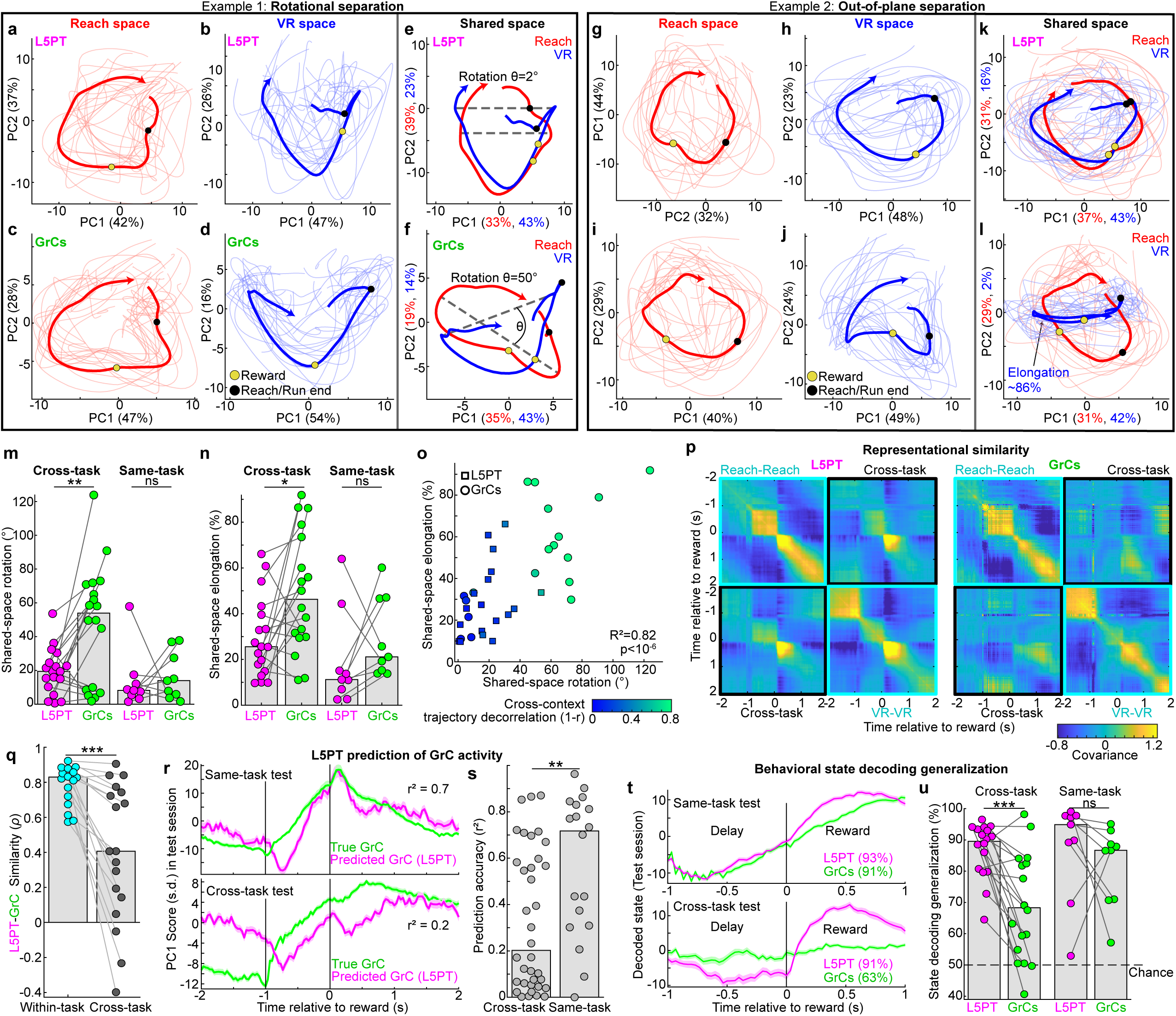
| GrC trajectories coherently reorient to separate contexts but preserve cortical geometry. **a–l,** Two types of cross-context GrC trajectory separation. **a–f,** Rotational separation. Top 2 PCs for L5 and GrCs in independent Reach **(a, c)** and VR **(b, d)** spaces. GrC trajectories preserved L5 geometry. **e,f,** Joint Reach-VR PCA (”shared space”). L5 trajectories aligned across tasks **(e)**, whereas GrC trajectories rotated 50° apart (**f**). **g–l,** “Out-of-plane” separation. **g–j,** Task space trajectories were similar across cell types (axes in **g** transposed for ease of comparison). **k,l,** In shared space, L5 trajectories aligned **(k)**, but the GrC VR trajectory “elongated” along PC1 (**l**, “out-of-plane” variance shifting). Thick/thin lines denote means and 15 trials closest to the mean. Single-trial trajectories omitted in **e,f** for clarity. Percentages denote variance explained. Arrowheads indicate epoch endpoint. Additional examples: **Extended Data Fig. 4**. **m,n,** Cross-task shared-space trajectory rotation angles (**m,** Procrustes, p=0.007) and elongation magnitudes (**n,** loss of circularity; p=0.02) were substantially larger for GrCs (same-task controls: p=0.6 and 0.3). **o,** Cross-context trajectory reorientation vs decorrelation (1-r mean across PCs 1-2). Rotation and elongation strongly predicted decorrelation (adjusted R^2^=0.82, p<10^-6^, F-test). **p,q,** Representational Similarity Analysis (RSA). **p**, Representative temporal covariance matrices. Within-task (cyan), GrC matrices preserved L5 geometry (Spearman’s ρ=0.88). Between tasks (black) GrC matrices were largely “blank” relative to L5 (ρ=0.05). **q,** Quantification (*ρ*) of L5-GrC RSA within versus across tasks (p=0.0003). **r,s,** L5 prediction of GrC activity (linear reconstruction of top 2 GrC PCs from 20 L5 shared-space PCs). **r,** Trial-averaged GrC PC1 activity overlaid with L5-predicted reconstruction for held-out test sessions (mean±s.e.m. across 143 and 120 trials). **s,** L5PT predicted GrC dynamics accurately in same-task tests but poorly in cross-task tests (p=0.008, two-sided Mann-Whitney U test; **Methods**). **t,u,** Behavioral state decoding. **t,** Trial-averaged LDA projection traces discriminating delay from reward (mean±s.e.m. across 140 and 78 trials). **u,** State decoding generalized better for L5PTs than GrCs across tasks (p=0.0009) but not for same-task controls (p=0.4). All cohort quantifications across 18 cross-task (9 mice) and 9 same-task (7 mice) session pairs. All paired comparisons used the Wilcoxon signed-rank test.

To determine the geometry of the “scrambled” cross-task GrC sequences, we next examined the shared-space projections. L5PT trajectories for VR and Reach remained tightly aligned, consistent with the principle of manifold reuse (**Fig. 4e,k**). However, the GrC trajectories coherently reoriented, revealing the underlying logic of contextual remapping. In Example 1, the GrC trajectories **rotated** apart in the shared plane, while preserving their intrinsic shapes (**Fig. 4f**). In Example 2, the VR trajectory **elongated** (**Fig. 4l**), indicating that variance rotated “out-of-plane” into orthogonal dimensions. These geometries demonstrated that the cross-context phase shifts and polarity inversions seen in individual GrCs (**Fig. 3d-f**) were in fact single-cell signatures of population-level rotations (additional examples in **Extended Data Fig. 4a-p**). Quantitatively, both “in-plane rotation” (Procrustes angle) and “out-of-plane” separation (elongation, i.e., loss of circularity, **Methods**) were substantially larger for GrCs than L5PT (**Fig. 4m,n**; rotation: 54±12° GrCs vs 19±3° L5PT; elongation: 46±9% GrCs vs 26±5% L5PT). Crucially, these effects were absent in cross-day same-task controls. Together, the magnitude of these geometric transformations predicted the functional outcome: cross-task trajectories with larger rotations exhibited greater temporal decorrelation (**Fig. 4o**, R^2^=0.82 multiple linear regression). Thus, rather than chaotically remap across tasks, GrC trajectories were coherently reoriented apart within largely overlapping subspaces.

### GrCs preserve within-task L5PT geometry

The PCA results revealed a tension: while GrC trajectories diverged between tasks, they appeared grossly similar to their L5PT counterparts within each task. To precisely test whether GrC trajectories preserved cortical representational geometry within each task, despite reorienting between tasks, we used Representational Similarity Analysis (RSA)^41^. RSA uses “second-order isomorphism”^42^ to compare the distances between timepoints in the full high-dimensional neural activity space, and is thus insensitive to differences in the neural ‘coordinate system.’ We computed similarity matrices for both tasks, organizing them into “within-task” versus “cross-task” blocks. Within each task, L5PT and GrC representational geometries were remarkably similar. For instance, in **Fig. 4p** Reach blocks, both cell types displayed a stable population state (high-similarity “square”) spanning the delay period [-1, 0] s, with a sharp transition after reward delivery. Similarly, in VR, both populations displayed high similarity throughout the running epoch. Thus, GrC trajectories robustly preserved within-task representational similarity structure of L5PT (**Fig. 4q**, Spearman’s *ρ*=0.83±0.03).

Between tasks, however L5PTs and GrCs diverged sharply. In **Fig. 4p**, L5PT displayed similarity *across* tasks, but the GrC cross-task block was dominated by dissimilarity, effectively “blanking out” L5PT’s cross-context generalization (**Fig. 4q**, *ρ*=0.41±0.2). Finally, cross-task separation did not come at the cost of within-task geometric distortion (p=0.4, **Extended Data Fig. 4q**). GrC representations thereby satisfy a dual objective: separating contexts by reorienting trajectories, while rigorously preserving the cortical representational geometry of each.

### L5-GrC mapping reconfigures across tasks

These results suggested that the L5→GrC transfer function is context-dependent. To quantify this, we used L5PT activity (20 shared-space PCs) to predict concurrent GrC activity (2 PCs). We trained a regression model for each session and tested its generalization on a held-out session of either the same task or the opposing task (**Fig. 4r**). Cross-day same-task L5-to-GrC prediction accuracy was high (**Fig. 4s**, r^2^=0.72±0.2), indicating a stable mapping. However, cross-task prediction accuracy collapsed (r^2^=0.2±0.1). Thus, the L5PT-to-GrC mapping reconfigures across tasks—a consequence of the rotated GrC trajectories.

### GrC behavioral readout remaps

This geometric reconfiguration dictates that a fixed downstream “readout” of behavioral state cannot generalize across tasks for GrCs, unlike the stable L5PT scaffold. To explicitly test this, we trained a decoder for each session using the top 3 shared-space PCs to distinguish two behaviorally distinct states conserved across both tasks: the delay ([-1, 0] s) versus the reward period ([0, 1] s). We then tested it on a held-out session from either the same task or the opposing task (**Fig. 4t**). Both L5PT and GrCs decoded behavioral state with high accuracy in cross-day same-task tests (95±6% L5PT, 87±6% GrCs, **Fig. 4u**). In cross-task tests, however, L5PT decoders remained strikingly accurate (90±2%), but GrC decoding accuracy collapsed (68±6%). This confirms that GrC representations effectively “encrypt” the cortical temporal scaffold into a context-specific format, preventing a static readout from generalizing across tasks.

### GrC task separation grows with learning

To investigate the relationship between learning and context separation in GrCs, we compared novice and trained mice. Although “Novice” data were acquired in consecutive Reach and VR sessions immediately following motor pre-training, animals at this stage lacked predictive knowledge of either task’s shared temporal structure. Most novice animals used a “reactive” licking strategy concentrated after reward delivery in both tasks, and shared-space PC1 activity for both L5PT and GrCs was typically strongly correlated across tasks (**Fig. 5a-c**). In trained animals, behavior converged on a “predictive” strategy with elevated licking before reward, and L5PT PC1 activity remained highly correlated across tasks (**Fig. 5d,e, Extended Data Fig. 5a-g**). Conversely, GrC PC1 remapped across tasks (e.g., “inverting polarity”; **Fig. 5f**). Quantitatively, GrC PC1 cross-task decorrelation significantly exceeded that of L5PT, especially in trained mice (**Fig. 5g,h**). However, “trained” performance varied: for example, some mice acquired predictive licking in only one task (**Extended Data Fig. 5h**). Thus, we next compared the amount of GrC context separation to *dual-task* proficiency (the minimum predictive licking score across both tasks). We found a strong positive relationship: the highest dual-task proficiency often accompanied the strongest GrC cross-context decorrelation (**Fig. 5i,j** *ρ*=0.6). This revealed a striking dissociation: as animals gained dual-task proficiency—resulting in *similar* predictive licking across tasks—their GrC representations became increasingly *distinct*.

**Figure 5.**
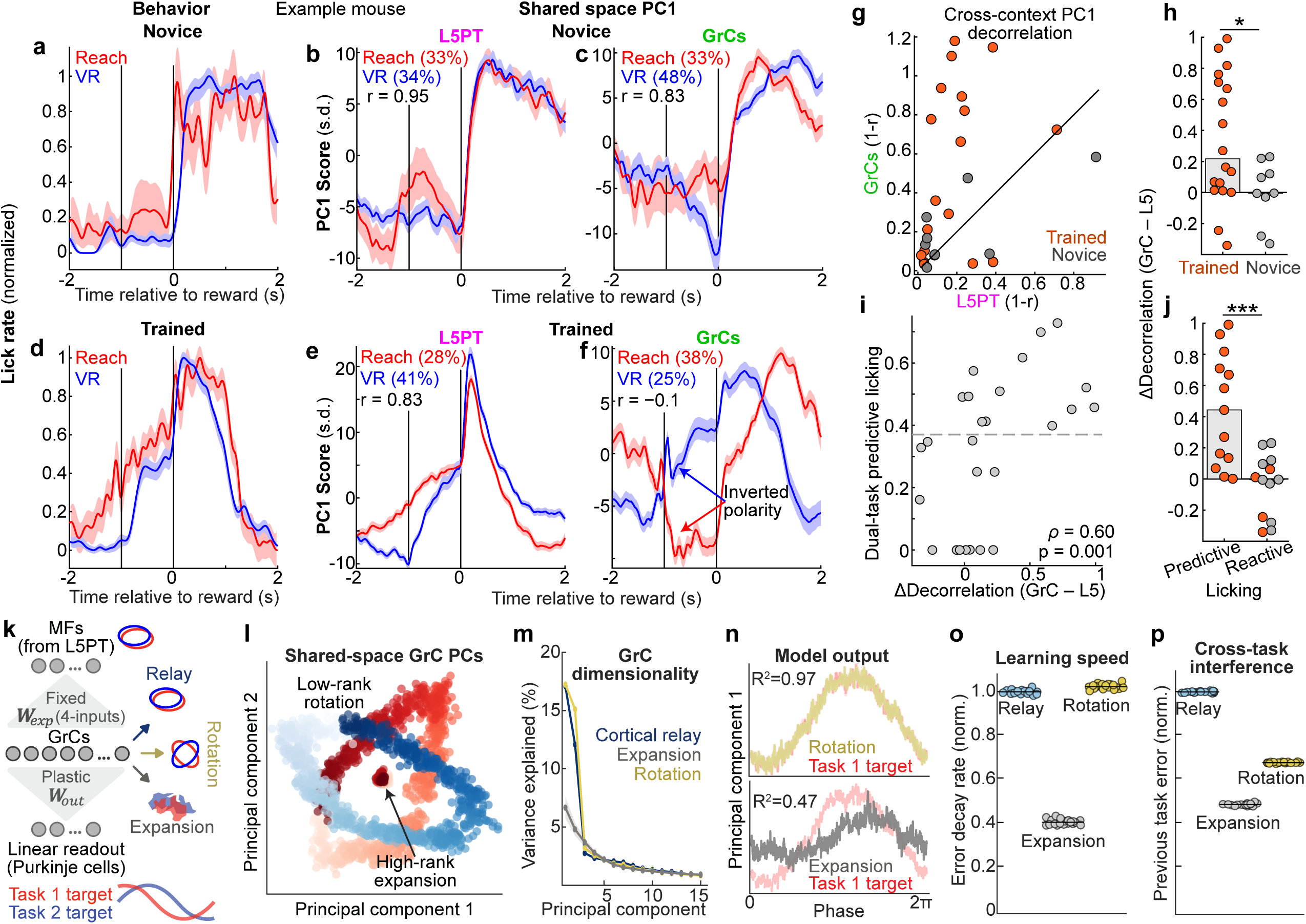
| Separating contexts via trajectory reorientation facilitates rapid dual-task learning. **a-f,** Representative Novice (**a-c,** consecutive sessions after pre-training) and Expert VR-Reach session pairs (**d-f**). Novice licking concentrated after reward **(a)**, and PC1 strongly correlated across tasks for both cell types (**b,c**; 15 Reach/46 VR trials). Expert licking became predictive **(d)**. Across tasks, while L5 PC1 activity remained correlated **(e)**, GrC PC1 **(f)** inverted polarity, despite PC1 explaining comparable variance in both cell types (percentages; 105 Reach/125 VR trials). Shaded regions show s.e.m. **g,h,** Comparison of L5 and GrC cross-context PC1 decorrelation (1-r). **g,** Scatter plot of Novice and Trained sessions. **h,** ΔDecorrelation (GrC PC1 minus L5 PC1) became modestly higher with training (p=0.037, two-sided Mann-Whitney U test, 18 Trained and 9 Novice session pairs). **i,j,** ΔDecorrelation (GrC minus L5) was higher in session pairs with better dual-task predictive licking (*ρ*=0.6, p=0.001; **Methods**). **j,** “Reactive” licking session pairs (including Novice and poor-performing Trained pairs) exhibited less cross-context GrC decorrelation than “Predictive” pairs (p=0.0009, two-sided Mann-Whitney U test, 26 pairs; Reactive/Predictive defined as below/above the dashed line in **i**). **k-p,** Simulated dual-task learning. **k**, Network architecture to predict kinematic targets via plastic linear readout at the Purkinje cell synapse (W*_out_*) from fixed GrC representations. Three GrC architectures were simulated: (1) Cortical relay; (2) High-rank expansion; and (3) Low-rank rotation. **l,** Latent GrC geometry. Rotation (open loops) orthogonalized the principal manifold axes while preserving structure; Expansion (center “dot”) collapsed manifolds into spectrally whitened representations. **m,** Effective dimensionality. Rotation and Relay models maintained low-rank input structure, whereas Expansion inflated dimensionality. Shaded regions show s.d. across 20 random network initializations. **n,** Output trajectories for Task 1 early in training (PC1 at Epoch 10). Rotation was more accurate than Expansion. **o,** Learning speed. Relay and Rotation models learned more rapidly than Expansion (**Extended Data Fig. 6**). **p,** Cross-task interference. Performance on the previously-trained task while training the opposing task (normalized MSE). The Relay model suffered from interference (catastrophic forgetting). The Rotation model mitigated interference comparably to the Expansion model (**o**,**p** normalized to Relay mean).

### Manifold rotation for optimal learning

We reasoned that neural architectures must navigate a fundamental tradeoff between learning efficiency and context independence. The aligned, low-rank structure of L5PT facilitates rapid learning by constraining the neural state space^3^, but invites interference between contexts. Conversely, the high-rank structure predicted by canonical expansion theory minimizes interference but exacerbates the “curse of dimensionality”. We hypothesized that the cross-context trajectory reorientations we observed in GrCs strike a “Goldilocks” balance between mitigating interference and accelerating learning using low-rank scaffolds.

To test this, we simulated alternating two-task learning in an anatomically constrained mossy fiber-GrC-Purkinje cell circuit with fixed mossy fiber-GrC projections (4 random mossy fiber inputs per GrC; 100-fold expansion) and a plastic linear readout. Input manifolds were overlapping ellipses, mimicking the L5PT data. The tasks required the networks to predict future states along each manifold, but with different time horizons for the two tasks (i.e., different rotational angles in the neural state space; **Extended Data Fig. 6a-d**). We compared three cerebellar-like architectures with differing GrC representations (**Fig. 5k-m**):

1. “Cortical Relay”: random projection that preserves input geometry.
2. “High-rank expansion”^5–8,32^: sparse coding that orthogonalizes patterns but sacrifices smoothness.
3. “Low-rank Rotation”: task-specific reorientation of the input geometry, modeled after our data.

The Relay and Rotation models learned the tasks significantly faster than the Expansion model (**Fig. 5n,o**): unlike discrete pattern classification tasks, continuous dynamic prediction tasks require networks to reconstruct *local distances between points* in the input manifolds within each context (mimicking a central Purkinje cell function^43,44^). Consequently, by disrupting Euclidean neighborhood relationships, the expansion model learned very slowly (and not merely due to sparse coding: learning speed similarly collapsed for dense-coding Relay models driven by manually tangled trajectories, **Extended Data Fig. 6k-o**). *Across* tasks, however, the Relay model suffered from interference (catastrophic forgetting of Task 1 while learning Task 2) due to manifold reuse. In contrast, the Rotation model reduced interference with efficacy comparable to the Expansion model (**Fig. 5p**). Thus, rotating trajectories between tasks solves the generalization-interference dilemma by separating contexts but preserving the representational smoothness required to rapidly learn dynamic predictions.

## Discussion

By simultaneously recording L5PT cortical output neurons and cerebellar GrC input layer neurons during parallel learning of two new skills, we tested the role of a universal mammalian pathway in resolving the generalization-separation tradeoff. Unlike canonical expansion architectures that disrupt low-dimensional manifolds, we found that GrCs instead faithfully preserved the low-rank neural structures and representational geometry present in the cortex. To separate contexts with overlapping cortical dynamics while preserving their representational geometry, GrC trajectories coherently reoriented between tasks—effectively “rotating” the structures apart. Crucially, GrC trajectories diverged most strongly across contexts in expert dual-task performers. This division of labor solves a fundamental control problem, allowing the brain to efficiently reuse generalized neural dynamics but still drive distinct policies.

High-rank expansion is optimal for *discrete pattern classification*^45^ (e.g., odors^19^; spatial environments^21^; visual objects^20^). However, cerebellar function is irreducibly *dynamic*, requiring continuous interpolation across time and space^43,44^. For Purkinje cells, which operate as nearly linear integrators^46^, continuous interpolation requires preserved Euclidean neighborhood relationships in GrC state space. Critically, high-rank expansion of a manifold violates this geometric constraint: by adding extrinsic curvature, it causes Euclidean and geodesic distances to diverge^47^, which exponentially increases the “covering number” of GrCs required to tile the manifold (the “curse of dimensionality”). Commensurately, such manifold “crumpling” would catastrophically increase the training samples required to support continuous generalization^33^. Here, we demonstrate that GrCs avoid this “maximal separation” strategy. Even the VR task, which required whole-body control during high-dimensional sensory flux, yielded dominant GrC subspaces of only ∼rank 4.

This raises the question: if rank remains low, why does the brain require an immense number of GrCs? While we investigated the separation of only two neural trajectories, the brain must manage hundreds of contexts—creating a “packing” problem. While the number of linearly separable patterns grows with high-rank expansion (the classical solution ascribed to the GrC layer), it also scales with the embedding dimension^4,32^— which is vastly larger for GrCs than any other circuit. Given the need to preserve smooth geometries for continuous generalization^2^, the circuit may instead rely on the sheer size of the GrC layer to separate many trajectories without resorting to “crumpling”. While “smoothness” is a theoretical interpretation of computational models rather than a directly measured quantity, the concept is strongly linked to the observed preservation of low-rank structure and representational geometry. Yet, prioritizing smoothness may impose another constraint: many GrCs were active simultaneously. If this “dense” code^37,38^ were “partitioned” by context^6–8,32^, it would rapidly saturate the circuit’s capacity. Instead, by maintaining overlapping active populations that reconfigure their *geometry* across contexts, the circuit mitigates interference while preserving representational density^39,48^. Consequently, while a single context exhibits a dense and low-rank code, the *collective* dimensionality of GrC representations can be high—fully utilizing the immense coding capacity.

A critical question concerns the synaptic mechanisms that could support reorienting trajectories. We first considered whether the observed L5-GrC divergence could simply reflect experimental recording bias rather than a bona fide representational shift. Evidence argued against *spatial* recording bias (i.e., recording from unconnected populations). First, RSA revealed that GrCs preserved the representational similarity structure of the cortex with high fidelity (*ρ*=0.83, **Fig. 4**). Second, CCA identified stable L5-GrC interaction weights with high predictive power across tasks (**Fig. 3**). Evidence similarly argued against *temporal* recording bias. While Ca^2+^ indicators act as temporal low-pass filters, our specific indicators actually permitted nearly 4-fold higher temporal resolution for GrCs than for L5PTs. Although this should reveal faster and thus higher-dimensional GrC signals, we found very little difference in effective dimensionality (**Fig. 2**). Finally, these findings were not driven by selection bias: within-task L5-GrC representational equivalence and cross-task GrC reorientations remained robust even when re-computed using all tracked cells, regardless of reliability (**Extended Data Fig. 7**). Instead, these data suggest that the L5-GrC divergence reflects the reorientation of shared, low-dimensional geometries. However, we note that equivalent representational geometry does not guarantee that the neural state is governed by the same underlying dynamical system^49^. Moreover, despite these premotor L5PT-GrC linkages, the large diversity of neocortical mossy fiber inputs to posterior cerebellar lobules leaves the precise locus of the cross-task GrC transformation—the GrCs alone or partially inherited from other pre-cerebellar sources that also remap between contexts—a critical question for future research.

Nevertheless, the GrC layer has distinctive anatomical advantages that could support reorienting representational geometries. Specifically, the GrC layer could perform gain-modulated integration of the cyclic cortical dynamics. Since low-dimensional geometries are widespread across the cortex, many GrCs must integrate multiple ‘in-manifold’ mossy fibers (especially given redundant input sampling^50^). “Remixing” these inputs—effectively rescaling the ‘sine’ versus ‘cosine’ cycle components—would directly produce the rotational signatures we observed in individual GrCs (**Fig. 3d-f**). Consequently, if input remixing were structured across the layer as a contextual ‘gain field’^51^, the entire population, and thus the entire cortical manifold, would undergo a coordinated transformation. Similarly, enhanced separation with learning (**Fig. 5**) might reflect emergence of a reliable ‘context’ multiplier. Mechanistically, GrCs would therefore operate as a tunable filter bank, where context scales the mixing coefficients that synthesize task-specific reorientations from a shared set of cortical primitives. This provides a compelling rationale for localizing this computation to the cerebellar cortex: because the GrC embedding space is uniquely high-dimensional, even randomly reorienting many low-dimensional manifolds in this way is likely to yield mutually orthogonal geometries^52^.

The extensive recurrent connectivity of the neocortex is anatomically suited for generating these stable, low-rank dynamical systems^53^. However, learning such manifolds is computationally expensive, requiring precise synaptic tuning to form a latent landscape^54^. This strongly incentivizes reusing dynamical ‘motifs’ across contexts to leverage shared underlying structure^55^, unless precluded by incompatible input-output mappings^13,18,56^. Yet, even when feasible, reuse entangles distinct neural trajectories^17,39^. The cerebellar architecture—defined by extreme numerical expansion—provides a complementary solution. By projecting cortical trajectories into a massive embedding space, the cerebellum can separate overlapping dynamics while preserving their representational geometry. Together these strategies yield an effective division of labor: the cortex specializes in slower learning^57^ of generalized dynamic primitives^58^, while the cerebellum provides the high embedding dimension needed to rapidly reconfigure these for contextually specific policies^26^. More broadly, this structural motif – using expansion layers to coherently reorient low-dimensional trajectories – provides a computational blueprint for continuous prediction and control in biological or engineered systems.

## Materials and Methods

### Mice

All dual-site imaging experiments used Math1-Cre×Ai93(TIGRE-LSL-TRE-GCaMP6f)×ztTA(R26-CAG-LSL-tTA) mice (aged 6–18 weeks), which express GCaMP6f selectively in cerebellar granule cells (GrCs; Math1-Cre: Jackson Labs 011104; Ai93: Jackson Labs 024103; ztTA: Jackson Labs 012266;, 10 mice total, 5 male). Optogenetic studies employed Gabra6-Cre×stGtACR1 mice (Gabra6-cre: MMRC 15966; LSL-stGtACR1: Jackson Labs 037380; 4 female mice). No statistical methods were used to predetermine sample sizes, but our cohort sizes are consistent with established standards in the field. Randomization of animals to experimental groups and investigator blinding were not relevant to this study, as all animals were in the same experimental group, all primary comparisons were within-subject across-task/cell type/learning stage, and data analysis pipelines were computationally automated. All procedures were approved by the NIH Animal Care and Use Committees.

### Viral Injections

An intersectional viral strategy was used to selectively target pons-projecting L5PT neurons in the premotor cortex. We injected AAVretro-EF1a-Flpo (Addgene 55637) into the basal pons and AAV1-Ef1a-fDIO-jRGECO1a (Addgene 128317) into the premotor cortex. Virus was injected at a titer of ∼10¹² genomes/mL.

Mice were anesthetized with isoflurane (1.5–2% in ∼1 L/min O₂) and mounted in a stereotaxic device (Kopf Instruments). A midline sagittal incision exposed the skull, which was cleared of connective tissue. Two ∼300 µm holes were drilled over the pons (0.3 and 0.8 mm right of midline, 3.9 mm posterior to bregma), and 1–2 holes were drilled over the premotor cortex (1.1–1.5 mm right of midline, 0.75–1.75 mm anterior to bregma, corresponding to the “rostral forelimb area” of the premotor cortex). Glass capillaries (∼25 µm tip diameter) delivered 300–400 nL of AAV-EF1a-Flpo at depths of 5.2 and 5.7 mm below the pial surface at the pontine sites, and 500 nL of AAV-Ef1a-fDIO-jRGECO1a at ∼800 µm depth at the cortical sites. Pipettes remained in place for 5 minutes before withdrawal. Expression was allowed to proceed for 4–5 weeks prior to imaging.

### Cranial Window and Headplate Implantation

Cranial windows were implanted 1–2 days after viral injections. To avoid mechanical collision between the cortical and cerebellar objectives, placement of the two windows was planned using computer-aided design (CAD) software.

A ∼10 mm x 8 mm patch of skin was excised over the skull. The exposed skull was cleared of soft tissue, and the skin incision edges were sealed with VetBond (3M). A primary cerebellar headplate (1.3 mm thick stainless steel, 5 mm opening) served as the main fixation device, while an auxiliary cortical headplate (1 mm thick, 2 screw holes) restricted frontal skull motion. A custom holder ensured consistent relative placement of the two headplates during implantation.

For the cortical window, a 3.5–4 mm craniotomy was centered 1.5 mm anterior to bregma and ∼1.5–2 mm right of midline. A 3 mm #0 coverslip glued to a stainless steel ring was positioned at a 20° azimuthal angle forward from the vertical plane and 25° clockwise from the sagittal plane.

For the cerebellar window, the craniotomy was centered over the post-lambda suture and ∼2 mm left of midline to access left lobules simplex, crus I, and crus II. The window assembly was depressed onto the brain at a 25° angle clockwise from the sagittal plane and a 50° azimuthal angle from the coronal plane. Windows were cemented with Metabond (Parkell), and headplates were cemented to the skull.

### Histology

Mice were deeply anesthetized with isoflurane, and then transcardially perfused with 1x PBS followed by 10% formalin. The brain was then dissected out and left overnight in 10% formalin. Following overnight fixation, the brain was transferred to a 30% sucrose solution in 1x PBS until the brain sank to the bottom of the solution. We then embedded brains in OCT Compound (Fisher HealthCare) prior to sectioning. Sagittal sections (50 um) were collected using a Microm HM550 cryostat into well plates containing 1x PBS.

We performed immunostaining on sagittal sections by first washing 3 x 10 minutes with 0.1% PBS-T, then blocking in 10% NGS in PBS-T. We then stained the sections using Rabbit anti-RFP (Abcam, 62341, 1:1000 dilution) and Chicken anti-GFP (Aves Labs, GFP-1010, 1:2500 dilution) for ∼48h, followed by 3 x 10 minutes washes with 0.1% PBS-T, followed by Alexa 488 goat anti-rabbit (Abcam, AB150077) and Alexa 647 Goat anti-Chicken (Abcam, AB150171, both 1:500 dilution) secondary antibody staining for ∼2 hours. Slides were imaged using a confocal microscope (Zeiss LSM 510) illuminated with the 488 and 647 nm lasers.

### Virtual Reality (VR) Setup

The custom VR apparatus^30^ consisted of an air-supported 8-inch polystyrene ball (SmoothFoam) and a hemispherical dome displaying a projection reflected via a convex mirror. To facilitate dual-task imaging on the same microscope, the apparatus was modular. The hemispherical dome, projector, and mirror were mounted on a moveable cart. The air-supported ball was mounted on a 12” x 12” breadboard along with head fixation bars, an axle restricting ball rotation to the forward/backward axis, a water reservoir, and solenoids for water delivery and airflow control.

Ball rotation was transformed into VR movement by an optical mouse (Logitech G502 Hero) mounted at the ball’s equator. The VR environment (a 1D track with patterned walls) was generated in MATLAB using ViRMEn software (Princeton). An Arduino UNO controlled the solenoids and generated a 1 ms pulse per ViRMEn iteration that was recorded by the data acquisition system which also sampled the two microscope frame clocks (PCI-6221, National Instruments, sampled at 5 kHz). The ViRMEn update rate was ∼10 ms.

To minimize light contamination, the projection was filtered (Kodak Deep Blue 47 Wratten Filter), and the red-channel PMT (cortex arm) was shielded with Cinefoil (Rosco). Additionally, a custom cover shielded the front lens of the cortical objective (Nikon 16x, 0.8 NA).

### VR Task Structure

Mice were teleported to the start of the track, and the air solenoid opened. After the mouse reached the reward zone (indicated by floor/wall pattern changes), the air solenoid closed, the projection froze, and a water reward was dispensed after a 1 s delay. Trials were aborted if the reward zone was not reached within 30 s. A 2 s inter-trial interval (black screen) followed every trial.

### Forelimb Reach Apparatus

We used a custom two-axis robotic manipulandum with modified data acquisition hardware and motor driver/encoder counting electronics. Device control was programmed in LabVIEW (National Instruments, NI) via a CompactRIO chassis (cRIO-9063) communicating with a Windows PC. A NI9401 digital I/O module generated PWM signals for the H-bridge motor drivers (Texas Instruments DRV8870EVM) and recorded the two microscope frame clock signals. Motor current was read via an ACS70331 sensor and sampled through a 9215 analog input module. DC motor rotation (Maxon DCX22) was measured by an optical encoder (Gurley R120) via a NI9411 differential digital input module.

The software utilized nested control loops: a 10 kHz FPGA loop for motor current and encoder readings; a 1 kHz real-time loop for geometric transformations, force calculations, and data buffering; and a high-level Windows PC loop for trial state management and data logging.

### Reach Task Structure

The task required a linear 8 mm reach. Mice self-initiated trials by pushing the handle. A reach >6 mm was scored as successful, triggering a water reward after a 1-s delay. Water was dispensed via a gavage needle equipped with a capacitive lick sensor. Trials terminated if movement ceased (>3 mm distance) for >100 ms. In all cases, the handle locked in place for 2-s after the end of the reach, and then automatically returned to the animal over the subsequent 2 s.

### Behavioral Training

Mice were water-restricted (maintained at >80% free-feeding weight) and monitored daily for health markers. Pre-training occurred for 7–10 days prior to imaging.

#### Pre-training

VR pre-training proceeded in stages: balancing and running short distances (∼20 mm) (3–5 days), followed by the full 60 mm track until performance reached ∼60 trials in 15 minutes (1–2 days). Subsequently, mice were pre-trained on the Reach task for 1–2 days until achieving pre-training proficiency comparable to the VR task.

#### Training and Recording Schedule

After pre-training, animals transitioned to alternating dual-task training for a median of 18 sessions (range: 11–22). Animals were trained on one task per session (1 session per day, typically across consecutive weekdays). Training generally alternated tasks across successive days (e.g., A-B-A-B); however, tasks were occasionally repeated on consecutive days if a mouse’s performance in one task required additional reinforcement to maintain dual-task balance.

Imaging was only performed on a subset of training days to minimize stress and accommodate the technical demands of dual-site alignment. Imaging data was grouped into three types:

- “Novice” cross-task session pairs were derived from consecutive training days immediately following pre-training (e.g., VR on Day N and Reach on Day N+1).
- “Trained” cross-task session-pairs were imaged at the end of the training period for all mice, with additional imaging midway through training for 5 of 9 mice. Sessions in Trained cross-task pairs were separated by 1 day (14 of 18 pairs), 2 days (2 of 18), or 5 days (2 of 18), contingent on successful cross-task dual-site optical alignment.
- “Same-task” control session-pairs were imaged only at the end of the training period; where alignment permitted, one session from these pairs was shared with the cross-task dataset to maximize data utility. Sessions in Same-task pairs were separated by 2 days (6 of 9 pairs) or 1 day (3 of 9).

Of the 10 mice in the study: one contributed Novice but not Trained data; one contributed Trained but not Novice data; and 2 contributed Trained cross-task but not same-task data due to optical alignment.

### Optogenetic Studies

Gabra6-Cre mice were crossed with LSL-stGtACR1 mice to generate double transgenics expressing soma-targeted GtACR1 inhibitory opsin in GrCs. Following window implantation and one week of single-task training, experts were subjected to perturbation. A ferrule-terminated optical fiber (200 µm core, 0.39 NA) was positioned ∼1 mm above the cerebellar window. A 594 nm laser (Coherent OBIS LX) delivered 5–15 mW (CW at fiber tip). During interleaved laser-on trials, a TTL signal triggered the inhibitory opsin in GrCs during the post-movement delay.

### Dual-site two-photon microscope

Building on a previous design^28^, we engineered a custom microscope with two independent, mechanically articulating arms to facilitate chronic dual-task imaging. We incorporated joystick-controlled (Zaber X-JOY3) 3D translation for both the mouse platform and the cortex imaging arm (”left arm”). The mouse positioning assembly utilized two long-travel, high-accuracy linear stages (Zaber X-LRQ150AP-DE51C) for X-Y translation and a high-load vertical stage (Zaber X-VSR40A, 40 mm travel) for Z translation, which supported the weight of both custom behavioral apparatuses via a custom adapter.

The independent cortex imaging arm was mounted on two heavy-duty linear stages (Zaber X-LRQ075HP-DE51, 75 mm travel) for X-Z translation. To accommodate the spatial constraints between the two objectives, Y translation was provided by a compact linear stage (Zaber LSA25A-T4A, 25 mm travel). Custom titanium adapters coupled the translation stages to the imaging assembly, while custom aluminum adapters mated the “floating” elliptic mirrors—required to periscope the beam across the translation stages—to Thorlabs ER rods for x-y-z positioning.

The cortex arm was equipped with a 16x 0.8 NA objective (Nikon CFI75 LWD 16X W) and excited using a 2W 1064 nm laser (Spark Alcor Dual) with a high-speed power modulator (Thorlabs OM6N). The cerebellum arm (”right arm”) was equipped with a 40x 0.8 NA objective (Olympus XLUMPlan) and excited using a 2W 920 nm laser (Spark Alcor Dual). Fine cross-day depth alignment employed objective z-Piezos (Thorlabs PFM450E) on both arms.

Data acquisition utilized the MBF vDAQ platform, running two parallel instances of ScanImage software (MBF Bioscience) on a single PC to independently control and acquire data from both imaging arms through a single vDAQ board. Both arms produced independent frame clock signals that were sampled by the behavioral acquisition hardware (either the Reach cRIO or the VR PCI-6221) to synchronize the two imaging streams with the behavioral data.

### Two-Photon Microscopy

Cortex imaging used the red-shifted indicator jRGECO1a excited at 1064 nm, focused to a depth of ∼600-800 μm, and with ∼70-140 mW of power at the objective. Cerebellar imaging used GCaMP6f excited at 920 nm, focused to a depth of ∼100-200 μm, and with ∼70-90 mW of power at the objective. Images (512x512 pixels) were acquired at 30 Hz.

### Cross-Rig and Longitudinal Field of View Registration

#### Microscope Calibration for Cross-Rig Transfer

To match the relative orientation of the two objectives across the different behavioral rigs (VR and Reach) utilizing the same microscope, a one-time structural calibration was performed using a test animal. On the VR setup, the optical axes of both the cerebellar and cortical imaging arms were aligned to their respective cranial windows via low-power visible laser back-reflection, and the pitch and yaw axes of both arms were fixed in place. The calibration animal was then immediately transferred to the Reach rig and placed under the microscope. There, the head-fixation posts were translated and rotated until both cranial windows achieved near-perfect back-reflection alignment with the previously fixed objective axes. This one-time calibration guaranteed that the relative angular geometry between the two objectives remained within ∼2 degrees on each axis when transferring experimental animals between rigs (even though absolute angles varied between animals by ∼±5 degrees).

#### Longitudinal Registration of Cerebellar Granule Cells (GrCs)

Due to expression density and small soma size, longitudinal and cross-task GrC tracking required precise three-dimensional coordinate mapping. On the initial imaging day, following data acquisition, the cortical objective was retracted, and the exact position of the three mouse translation stages was recorded via Zaber Launcher software. Under a wide-field camera, we used the mouse stages to translate the field of view to a distinct anatomical landmark (e.g., a major blood vessel intersection) to capture a reference image, and the relative 3D translation vector between this landmark and the GrC imaging site was calculated. On subsequent imaging days (or during transfer to the alternative task rig), the optical axis was first aligned via back-reflection. The translation stages were then used to visually locate the anatomical landmark by matching the live wide-field camera feed to the prior session’s reference image. The previously calculated 3D mouse translation vector was then applied in Zaber Launcher to physically return to the precise imaging site (typically within 100 μm). Final alignments were achieved during live two-photon imaging by matching the real-time structural features to the mean reference two-photon image from the prior session using both the mouse stages and the objective z-piezo.

#### Longitudinal Registration of Cortical Layer 5 Pyramidal Tract Neurons (L5PT)

Longitudinal registration of the L5PT population was performed after fine-positioning the cerebellar imaging field, and was achieved via direct visual alignment facilitated by the comparatively sparse viral expression, large somatic size, and restricted labeling area. Following optical axis alignment via back-reflection, the live two-photon field of view was manually matched to the previous session’s mean reference image by adjusting the positions of cortex arm’s isolated motorized stages and z-piezo depth until the somatic topology was visually confirmed.

#### Image Preprocessing

Data were motion-corrected using NoRMCorre (Simons Foundation) sequential large-displacement rigid followed by small-displacement non-rigid correction). Slow drifts were corrected by dividing out exponential fits to the frame-averaged fluorescence. Source extraction for GrCs and L5PTs was performed using cNMF (Simons Foundation), followed by manual curation. Traces were obtained by back-applying spatial filters and z-scoring.

#### Cross day registration and cell filtering

To track neurons across imaging sessions, we first computed image registration parameters between the mean intensity projection images of each session. These parameters were applied to transform the cell spatial filters from one session into the coordinate space of the other. Cells were initially matched automatically by thresholding the percent spatial overlap between the projected filters and the local filters.

To recover cells that were active in both sessions but missed by the automated sorting in one, we performed a “rescue” step. “Unmatched” filters from one session were projected onto the movie data of the other session. These candidates were then manually curated by verifying the presence of a visible “cell” in the z-scored movie (z-scored per pixel across all timepoints) at the times of their fluorescence peak activity timepoints. This procedure was performed bidirectionally (Session 1 onto Session 2, and vice versa). The union of these operations yielded a consensus set of neurons with validated activity in both sessions.

We subsequently filtered this population for response reliability. For each session, we calculated the split-half reliability as the Pearson correlation between the average activity on odd versus even trials, *r_odd:even_*. This raw correlation was corrected using the Spearman-Brown prediction formula: *R_raw_*=2\**r* / (1+*r*). To account for differences in trial counts across sessions, we standardized the reliability to a fixed count of 100 trials using the Spearman-Brown prophecy formula: *R_adj_*_=_*K*\**R_raw_* / (1+*R_raw_*\*(*K*-1)), where *K*=100/*N*_trials_. Unless otherwise indicated (e.g., **Fig. 2L-N)**, all downstream analyses were restricted to neurons that were jointly reliable (*R_adj_*> 0.4) in both sessions of the pair. To ensure this reliability filter did not inadvertently mask a large population of task-specific cells, we quantified “task-specific” prevalences using a conservative reliability criterion if in one task R_adj_>0.6 but in the other R_adj_<0.3. In cross-task comparisons, a near-identical 10% of L5PTs and 9% of GrCs were classified as task-specific. These rates were only marginally higher than in same-task cross-day control comparisons (8% of L5PTs and 7% of GrCs). This confirms that reliability-filtering primarily removed cells subject to condition-independent noise rather than biological task-specificity.

## Quantification and Statistical Analysis

All custom data analysis was performed in MATLAB (Mathworks).

Unless otherwise noted, quantifications in the text are reported as median ± standard error of the median (s.e._median_), computed as the standard deviation of the median across 10,000 bootstraps of the data. Unpaired comparisons used the Mann-Whitney U-test; paired comparisons used the Wilcoxon signed-rank test.

To compare the cumulative distributions of peak activity times between L5PT and GrC populations while accounting for session-to-session nested variance (**Fig. 2e,f**), we employed a custom permutation-based Kolmogorov-Smirnov test. The observed test statistic was defined as the maximum absolute difference between the session-averaged cumulative distribution functions (CDFs) of the two cell types. A null distribution was generated by randomly shuffling the session labels across both cell types 1,000 times, computing the maximum absolute difference between the mean CDFs of the pseudo-groups for each iteration. The p-value was defined as the fraction of permutations where the null statistic met or exceeded the observed statistic.

### Behavioral Analysis

Lick rates were calculated by binning lick sensor events at 1 kHz and smoothing with a Gaussian kernel (σ = 20 ms). Lick rates were normalized within each session using the session’s rewarded trial-averaged lick-rate trace: (lick rate - min(avg)) / (max(avg) - min(avg)). “Avg” was defined as the 1-s Gaussian-smoothed lick-rate trace over [-3, 2] s, excluding the first 0.2 s after reward delivery. Anticipatory licking quantifications (**Fig. 1t** and **Extended Data Fig. 2**) used the same normalized rates, with values below the baseline minimum, i.e. normalized values < 0, clipped to 0. The **”Predictive Licking Score”** (**Fig. 5** and **Extended Data Fig. 5**) was calculated to quantify the shift from reactive to anticipatory licking. For each task, the score was defined as the ratio of pre-reward licking to total licking: Lick_pre / (Lick_pre + Lick_post), where Lick_pre is the median across trials of the rate in the window [-0.5, 0] s relative to reward, and Lick_post is the median across trials of the rate from [1.0, 1.5] s. Trials were excluded from licking analysis if the capacitive sensor malfunctioned (became “stuck” high across any 1.5 s interval in the trial). The “dual-task predictive licking” score (**Fig. 5**) was defined as the minimum predictive licking score across the two tasks in each session pair.

Note: All analyses based on licking excluded 1 Trained session-pair due to a malfunctioning (“stuck”) capacitive sensor in the VR session. Thus, these panels have only 17 Trained session-pairs (**Fig. 1t**, **5i, 5j, Extended Data Fig. 5h, 7i,j**).

For visualization only, cohort (**Fig. 1**) and exemplar (**Fig. 5, Extended Data Fig. 5**) lick traces were Gaussian smoothed with σ=40 and 60 ms respectively.

### Single-Neuron Response Metrics

Peak activity times were determined from the maximum of the trial-averaged fluorescence trace [-2, 2] s relative to reward, with trial-averaged traces z-scored per-cell. The temporal width of the response was quantified as the Full Width at Half Maximum (FWHM), calculated as the time difference between the half-peak amplitude crossings of the trial-averaged trace, after Gaussian smoothing with *σ*=33 ms.

To estimate the true correlation of neural activity across two days (*r_true_*), we applied Spearman’s attenuation correction to the raw Pearson correlation (*r_raw_*) of the trial-averaged traces, accounting for the internal reliability (*r_xx_* and *r_yy_*) of the neuron in each context: *r_true_* = *r_raw_* / sqrt(*r_xx_* * *r_yy_*), where *r_xx_* and *r_yy_* are the split-half reliabilities defined above (odd vs. even trials) for the two sessions.

### Dimensionality analysis

For the dimensionality analysis only (**Fig. 2l-n and Extended Data Fig. 3a**), each session was considered independently. Rather than matching cells across days, all cells that were present and cleared the reliability criterion in each session were included. This maximized the number of available cells to best estimate the participation ratio (PR) and saturation curves. For the trial-averaged dimensionality analysis (**Fig. 2**), Principal Component Analysis (PCA) was performed on the matrix of trial-averaged activity in the window [-2, 2] s relative to reward delivery (Time × Neurons), after z-scoring each cell’s trial-averaged trace. The effective rank (dimensionality) was estimated using the Participation Ratio (PR): PR = (∑ *λ_i_*)^2^⁄∑ *λ_i_*^2^), where lambda are the eigenvalues of the covariance matrix. To assess the robustness of this metric to population size, we computed PR saturation curves by randomly subsampling the population at 20 intervals (from 1 neuron to N) and averaging the PR over 50 bootstrap iterations per interval. For the single-trial dimensionality analysis (**Extended Data Fig. 3a**), to reduce PR inflation due to uncorrelated optical noise, single-trial data were first Gaussian smoothed with *σ*=0.2 s.

### Task-space and shared-space low-dimensional dynamics

For the analysis of low-dimensional geometry across contexts (cross-task pairs or same-task controls across days, **Figs. 3–5 and Extended Data Figs. 3–5**), each session pair provided two matrices of trial-averaged activity, denoted as X_1_ and X_2_. Both matrices were constructed with dimensions of T×N_matched-cells_), where T represents the time window [-2, 2] s relative to reward. Prior to dimensionality reduction, each cell’s trial-averaged trace was z-scored and Gaussian smoothed (*σ=*20 ms).

Task-space PCA (or the same-task control equivalent) was computed directly on each individual matrix.

To identify neural modes shared across contexts (or days for same-task), we performed joint-task PCA. The paired trial-averaged matrices were concatenated along the temporal dimension to form a joint matrix X_Joint_ = [X_1_, X_2_]. PCA was applied to X_Joint_ to yield a common set of eigenvectors (joint PCs), and the activity for each session was subsequently projected onto these joint axes.

For example trajectory display only (**Fig. 4a-l, Extended Data Fig. 4a-l**), mean and single trial trajectories were Gaussian smoothed with σ=0.1 and 0.2 s, respectively.

### Canonical Correlation Analysis (CCA)

CCA was used to identify the “communicating subspace” between L5PT and GrC populations. We applied CCA to the concatenated whole session activity matrices for all cells in both tasks to find pairs of canonical variables (CVs) that maximized the correlation between L5PT and GrC activity. We then retained all CVs with r_L5-GrC_>0.1, and computed the L5-GrC correlation for each component in each task separately (r_L5-GrC_^Reach^ and r_L5-GrC_^VR^). Finally, to quantify the resulting cross-task stability of the communicating subspaces, we computed the correlation between r_L5-GrC_^Reach^ and r_L5-GrC_^VR^ across all retained CVs (**Fig. 3x**, r = 0.69).

### Geometric Analysis

- **Representational Similarity Analysis (RSA):** We quantified the preservation of representational geometry using temporal autocovariance matrices. Using the concatenated joint activity matrix X_Joint_=[X_1_,X_2_] defined above, we computed a 2T×2T similarity matrix S where each element *S_i,j_* represents the dot product of the population activity vectors at timepoints *i* and *j* (i.e., the unnormalized covariance). This metric was chosen to preserve information regarding the magnitude of population activation at each timepoint. To compare L5PT and GrC geometries, we calculated separate Spearman rank correlations (*ρ*) between the L5PT and GrC similarity matrices for the “diagonal” (within-context) and “off-diagonal” (cross-context) blocks, after vectorizing each block.
- **Procrustes Analysis:** To quantify the geometric rotation of the low-dimensional trajectories between tasks, we aligned the top 2 PC trajectories of the Reach and VR tasks using Procrustes superimposition (allowing translation, scaling, and rotation). The “Shared-space rotation” was the angular component of this transformation.
- **Circularity (isoperimetric quotient):** To quantify the geometry of neural trajectories, we calculated the Isoperimetric Quotient (often termed Circularity or Form Factor). This was defined as: *Circularity* = 4π\**Area*/*Perimiter*^2^, where *Area* is the area enclosed by the trajectory (calculated via the MATLAB polyshape function, which automatically resolves and merges any self-intersecting trajectory loops) and *Perimeter* is the total arc length of the trajectory (sum of Euclidean distances between consecutive timepoints). A value of 1.0 indicates a perfect circle, while values approaching 0 indicate increasingly elongated or convoluted trajectories.
- **Elongation (Loss of Circularity):** To quantify the “out-of-plane” warping of trajectories in the shared subspace, we defined Elongation as the reduction in circularity relative to the independent task space: *Elongation* = 1 - (*Circ_shared_* / *Circ_task_*). The summary quantification for each session pair was the maximum of this value across the two paired sessions.

### Decoding and Regression Models

- **L5** → **GrC Prediction:** We trained linear regression models to predict the temporal activity of each of the top 2 GrC shared-space PCs using the top 20 L5PT shared-space PCs as predictors. Models were trained on data from one session and tested on held-out sessions (either “same-task” or “cross-task”). Prediction accuracy was quantified as the squared Pearson correlation (*r^2^*) between the predicted and actual GrC PC scores. A model was trained for each “direction” (VR→Reach or Reach→VR for cross-task; Day 1→Day 2 or Day 2→Day 1) and we reported the bidirectional mean (**Fig. 4s**) for each of the top 2 GrC components (yielding 36 cross-task and 18 same-task datapoints from 18 and 9 session-pairs respectively).
- **Behavioral State Decoding:** We trained Linear Discriminant Analysis (LDA) classifiers to distinguish “Delay” ([-1, 0] s) from “Reward” ([0, 1] s) epochs. Input features for training were computed by averaging the activity across the duration of each epoch for every trial for each of the top 3 shared-space PCs. To visualize the decoding time course (**Fig. 4t**), we projected the continuous, time-varying activity of the test session onto the discriminant axis identified by the training session. Generalization accuracy (**Fig. 4u**) was defined as the percentage of correctly classified trial epochs in the test session.

## Neural Network Simulation

Simulations were performed in Python.

### Task geometry

We generated supervised training pairs (**x**, **y**) from circular manifolds with intrinsic dimension 1, embedded in a 100-dimensional ambient space. The intrinsic circle was defined as:

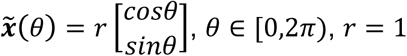

To better mirror neural data, we introduced anisotropy using a transform in intrinsic coordinates:

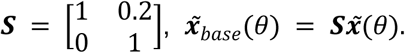

The resulting 2D manifolds were embedded into ℝ¹⁰⁰ using a random orthonormal basis ***E*** ∈ ℝ¹⁰⁰^×^², ***E***ᵀ***E*** = ***I***₂. We then defined two task-specific input manifolds that shared the same base circle, the same shear, and the same embedding plane, differing only by a translation perpendicular to the embedding plane:

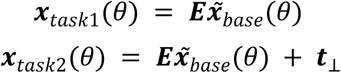

where ***t***_⊥_ ∈ ℝ¹⁰⁰ is constrained to be orthogonal to the embedding plane (***E***ᵀ***t***_⊥_ = 0). For each task, we sampled 1000 phases *θ*ᵢ ∼ *U*[0, 2*π*) and added pink noise to obtain the inputs:

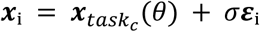

where ***ε***ᵢ is pink noise (*β* = 1), and *σ* = 0.15 and *c* corresponds to task identity (*c* ∈ [1, 2]). To construct the targets, we rotated the same base manifold in intrinsic coordinates using task-specific in-plane rotations:

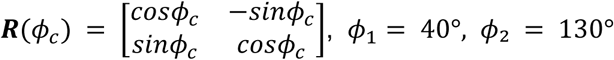

The noiseless target manifolds were (**Extended Data Fig. 6a-c**):

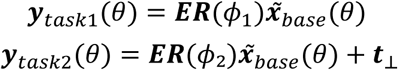

We then sampled these manifolds with pink noise again to obtain the targets ***y***ᵢ = ***y****_taskc_* (*θ*ᵢ) + *σ**ε***ᵢ, with ***ε***ᵢ pink noise (*β* = 1) of *σ* = 0.15. Thus, the dataset consists of paired samples (***x***ᵢ, ***y***ᵢ), where inputs come from unrotated task manifolds and targets from rotated task manifolds, while task separation is preserved by the same perpendicular translation term ***t***_⊥_.

The curriculum consisted of four sequential training phases (100 epochs each): task 1, task 2, task 1, task 2, implementing a continual-learning protocol (**Extended Data Fig. 6d**). Data were presented in shuffled mini-batches of size *n* = 32 within each phase.

### Model architectures

We compared three cerebellar-inspired neural network architectures designed to investigate how different computational motifs affect continual learning and interference. All models shared a common three-layer structure: an input layer receiving mossy-fiber patterns (**x** ∈ ℝ¹⁰⁰), an intermediate granule-cell layer (**h** ∈ ℝ¹⁰⁰⁰⁰), and a readout layer producing predictions (**ŷ** ∈ ℝ¹⁰⁰). For all models, the readout mapping was identical:

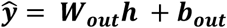

Crucially, to mimic the anatomical constraints of the cerebellum, the input connectivity was identical for all three models: the projection matrix ***W_exp_*** ∈ ℝ¹⁰⁰⁰⁰^×^¹⁰⁰ was fixed (untrained) and sparse, where each granule cell received input from exactly 4 randomly selected mossy fibers (4:1 convergence). The models differed only in how the GrC activation *h* was computed from this sparse input.

1. **Relay model.** The relay model implemented the standard 4-input sparse projection followed by ReLU, but without the normalization or high thresholds that sacrifice smoothness by disrupting Euclidean neighborhood relationships. This preserved the manifold geometry in the high-dimensional embedding:

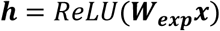

2. **High-rank expansion model.** The expansion model implemented the canonical Albus/Marr sparse coding theory. It used the same 4-input sparse projection but incorporated L2 input normalization (Golgi cell-mediated feedback^8^) and a fixed negative bias to enforce high population sparsity (∼1%):

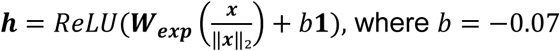

3. **Rotation model.** This model implemented the interference-reducing geometry observed in our data by applying task-dependent geometric transformations *before* the same 4-input sparse projection:

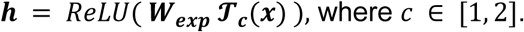

The input was decomposed into in-plane and out-of-plane components relative to the embedding plane *E*. Transformations (rotation and shear) were applied only to the in-plane component in intrinsic 2D coordinates, then recombined with the preserved out-of-plane component:

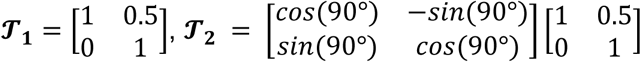

where task 1 applied only shear and task 2 combined a 90° rotation with shear. The full transformation can be written as:

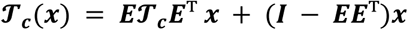

All linear weights were initialized with Kaiming normal initialization. Sparse masking was applied after initialization to enforce 4-input connectivity per granule cell. Only readout parameters (***W_out_*, *b_out_***) were trained using Adam with learning rate *η* = 0.0001 and mean-squared error loss:

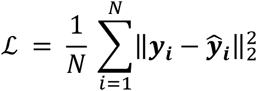

Training was repeated across 20 random seeds for each model type

### Model comparison metrics

#### Variance explained by principal components

Principal component analysis was performed on granule cell activations (combined across both tasks) to quantify the dimensionality of the population code. For each model architecture and random seed, we computed the proportion of variance explained by the first 20 principal components. Results are reported as mean ± standard deviation across random seeds.

#### Learning speed

Learning speed was quantified by fitting an exponential decay function to the loss trajectory: *L*(*t*) = *A* · *exp*(−*t*/*τ*) + *B*, where *t* is the epoch number, *A* is the initial decay amplitude, *τ* is the time constant, and *B* is the asymptotic loss. Learning speed was defined as 1/*τ*, representing the rate of exponential loss decrease (higher values indicate faster learning). To disambiguate learning speed from interference effects, speeds were reported for the first phase only. Fits were performed using nonlinear least squares optimization (scipy.optimize.curve_fit).

#### Previous task loss

To monitor catastrophic forgetting during continual learning, we tracked the mean squared error on the previous task’s data whilst training on the current task. For each training batch on task 2, we computed the loss on task 1 data using the current model parameters without gradient updates (and vice versa).

#### Granule cell overlap (Jaccard similarity)

To quantify the degree of shared neural representations between tasks, we computed the Jaccard similarity of active granule cells. For each sample, we binarized granule cell activations and calculated the Jaccard index as |*A* ∩ *B*| / |*A* ∪ *B*|, where *A* and *B* are the sets of active cells for tasks 1 and 2, respectively. We compared activations from models trained to completion on each task (checkpoint after phase 1 for task 1; checkpoint after phase 2 for task 2), then averaged Jaccard scores across all samples and compared random seeds across model types.

### Trajectory tangling control experiment

To test whether the geometric complexity of the input manifold accounts for differences in learning speed and cross-task interference, we conducted a trajectory tangling experiment. A sinusoidal transform was applied to the input manifold prior to all network layers, progressively distorting its smooth circular structure in the ambient space while preserving the norm of each point. The transform consists of *I* folds, where each fold is a single sinusoidal displacement in a random ambient-space direction that oscillates around the ring. Summing *I* folds with independent directions, frequencies, and phase offsets produces a complex, high-frequency “crumpling” of the manifold geometry. Specifically, for each fold i, a random unit vector ***d****_i_* ∈ ℝ¹⁰⁰ was drawn from the full ambient space, an integer-valued random frequency *f_i_* ∼ *U*(2, *I*) and phase offset *φ_i_* ∼ *U*(0, 2*π*) were sampled, and the displacement

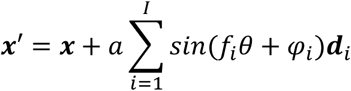

was applied, where *a* is the displacement amplitude and *θ* is the angular position of each point projected onto the manifold’s embedding plane. The result was then rescaled to the original norm. Increasing *I* increases the geometric complexity of the manifold as seen by the readout layer, while leaving the task structure and all network parameters unchanged (for visualisation, *I* was normalised to a tangling strength *s* ∈ [0, 1] across the tested range. This transform was applied to the relay model’s input across a range of tangling levels, and compared against the expansion model baseline (no tangling). If input geometry were the driver of slow learning and increased interference in the expansion model, tangling the relay’s input trajectory should push its behavior progressively toward the expansion model.

## Data availability statement

The data in this study have been publicly deposited in Dryad at 10.5061/dryad.9p8cz8x0p.

## Code availability statement

Code is publicly available at Zenodo (10.5281/zenodo.21461929.)

## Acknowledgements

We thank National Institute of Mental Health Section for Instrumentation for fabrication, the National Eye Institute Ocular Gene Therapy Core for viral packaging, and the National Eye Institute Building 49 Central Animal Facility staff for animal care and husbandry, and Yi Gu and Chris McBain and their lab members for designs for the VR navigation apparatus. We thank L Luo, and members of the Neocortex-cerebellum circuitry unit for helpful comments on the manuscript.

## Author contributions

M.G.G. and M.J.W. conceived the study. M.G.G. performed all *in vivo* experiments. M.G.G. designed and implemented the behavioral apparatuses with assistance from O.A. and M.J.W. M.G.G. performed surgeries and behavioral training with assistance from A.O., S.T., and M.J.W.. A.O. and L.R. assisted with histology. M.J.W. assisted with microscopy design. S.T. and L.D. contributed to data processing pipelines. M.G.G., L.D., and M.J.W. performed the neural data analysis and geometric quantifications. M.Wójcik and R.P.C. designed and implemented the neural network simulations. M.J.W. supervised the study. M.G.G. and M.J.W. wrote the manuscript with input from all authors.

## Ethics statement

This study complied with all relevant ethical regulations for animal testing and research. All animal procedures and experiments were performed in accordance with the guidelines of the National Institutes of Health and were approved by the Institutional Animal Care and Use Committee (IACUC) of the National Institute of Neurological Disorders and Stroke (NINDS).

## Additional Information

Correspondence and requests for materials should be addressed to Mark J. Wagner or Martha G. Garcia-Garcia.

## Extended Data Figures

**Extended Data Figure 1.**
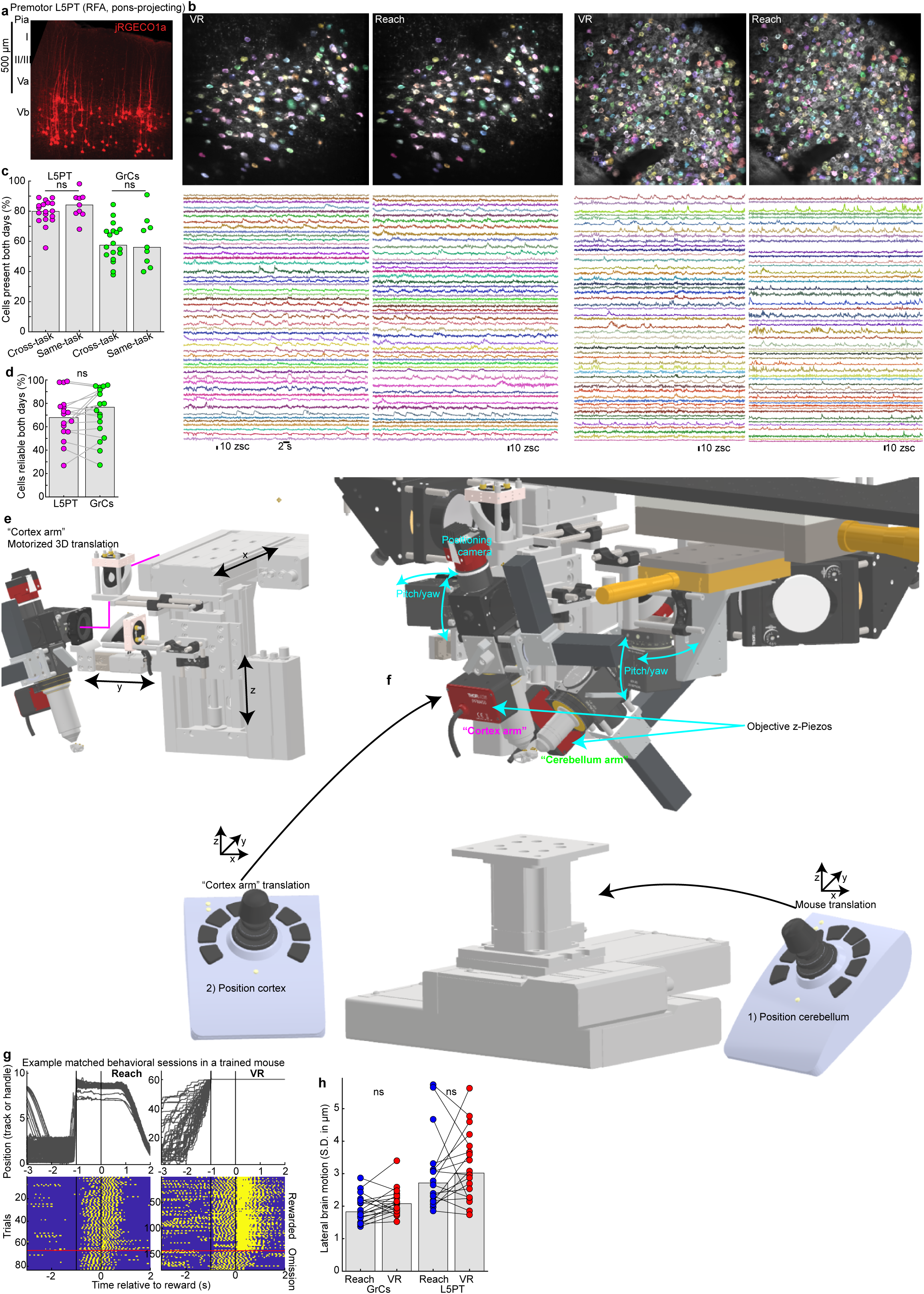
| Dual-task cortico-cerebellar two-photon imaging. **a,**Histological image of jRGECO1a expression in pons-projecting premotor cortical neurons, which are enriched in the deeper part of Layer V ∼700 µm below the surface of the brain. “RFA”, rostral forelimb area of the premotor cortex. **b,** For the example VR-Reach matched imaging session pair in Fig. 1, mean two-photon images show spatial filters for all L5PTs and GrCs with detected activity in both tasks (138 L5PT and 368 GrCs). Traces show 50 example neurons from each imaging session and cell type with colors corresponding to the cell maps above. **c,** Cross-day cell tracking retention rates. For each session pair, dots indicate the fraction of cells in session 1 that also had detected activity in session 2. While overall retention was lower for GrCs than L5PT due to optical constraints, there was no significant difference between cross-task and same-task retention rates within either cell type (L5PT: p=0.5; GrCs: p=1, two-sided Mann-Whitney U-test), demonstrating that tracking attrition was not driven by context-switching (18 cross-task and 9 same-task session pairs from 9 mice). **d,** Bars show fractions of cells in each cross-task session pair that were reliable across both tasks, which did not differ by cell type (p=0.3, Wilcoxon signed-rank test; 18 cross-task session pairs). **e,f,** Custom dual-site microscope mechanics enabling cross-day registration. To accommodate the challenges of tracking dual-region brain activity while alternating between two distinct behavioral apparatuses, we redesigned a dual-site two-photon microscope^28^ to equip both the “cortex arm” and the mouse platform with joystick-controlled motorized 3D translation. **e,** Isolated view of the redesigned “cortex arm”. Two high-load, long-travel (75 mm), high accuracy Zaber stages provided motorized “x” and “z” control (X-LRQ075HP-DE51). A third compact motorized Zaber stage provided 25 mm of “y” travel (LSA25A). **f,** Overall view of both the motorized cortex arm and the cerebellum arm with 16x 0.8 NA Nikon and 40x 0.8 NA Olympus objectives positioned over the imaging sites on a model mouse skull. Below, a motorized 3D translation platform with 150 mm x/y travel (Zaber X-LRQ150AP-DE51C) and 40 mm z travel (Zaber X-VSR40A) allowed positioning the cerebellum under the cerebellar imaging objective for both behavioral apparatuses. Registration followed a four-step sequence: (1) Align the cerebellar objective optical axis to the window using the “cerebellar arm” pitch/yaw rotational axes. (2) Position the cerebellar imaging site using the motorized translation platform; fine-tune depth via objective z-piezo; (3) Align the cortical objective optical axis using the left arm pitch/yaw rotational axes; (4) Position the cortex imaging site using the motorized cortex arm; fine-tune depth via objective z-piezo. **g,** Single-trial behavioral data from a representative matched Reach-VR session pair. Traces (top) show single position trajectories and Rasters (bottom) show binary lick sensor contacts with trials grouped into rewarded and omitted reward blocks for ease of visualization. **h,** Brain motion. Dots show sessions, quantified as standard deviation across all frames of the lateral motion correction computed by the image registration algorithm. Brain motion did not differ between tasks, but was slightly higher for L5PT likely because the primary skull fixation plate encased the cerebellar window while the cortical window was stabilized by a smaller auxiliary plate (GrCs: p=0.08; L5PT: p=0.3, Wilcoxon signed-rank test; 18 cross-task session pairs).

**Extended Data Figure 2.**
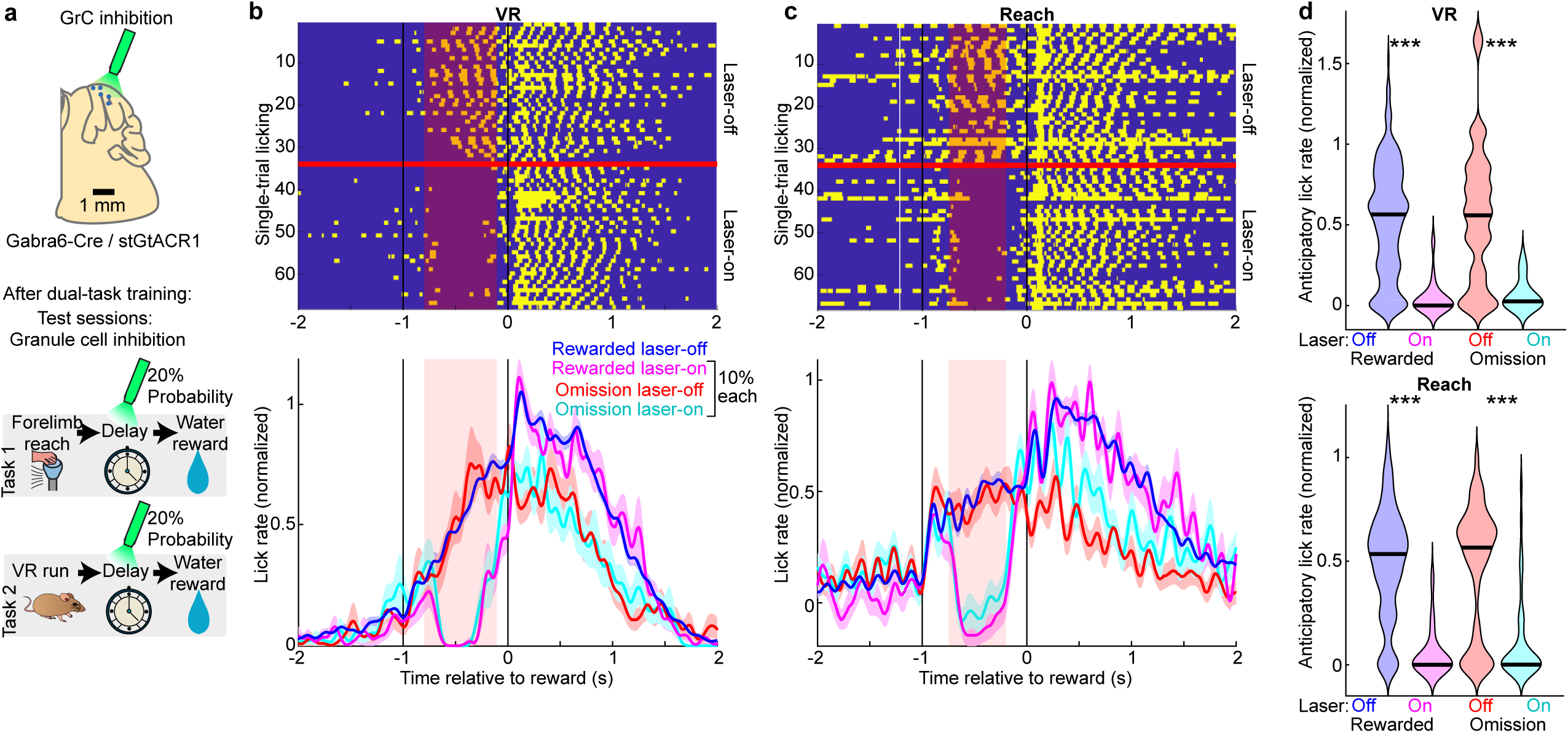
| GrC activity is necessary for the expression of anticipatory licking in both tasks. **a,**Schematic of the optogenetic paradigm for inhibiting GrCs during the delay period on interleaved perturbation trials (4 sessions from 4 mice). 70% of trials were rewarded laser-off; 10% each were: rewarded laser-on; omission laser-off; and omission laser-on. Posterior cerebellum was illuminated through a cranial window with a 594 nm laser in Gabra6-Cre×stGtACR1 mice, which express the soma-targeted inhibitory opsin GtACR1 in all GrCs. **b,c,** Licking behavior in the VR task **(b)** and Reach task **(c)** during control trials and randomly interleaved GrC inhibition trials (20%). Rasters (top) show binary lick contacts for rewarded laser-off and rewarded laser-on conditions (all 35 (VR) or 34 (Reach) laser-on rewarded trials and a randomly selected subset of laser-off rewarded trials drawn from 243 (VR) or 263 (Reach) trials from 4 sessions per task from 4 mice). Traces (bottom) show smoothed lick rates for all four conditions (normalized identically to the main dataset, using the rewarded laser-off data; Reach/VR trials: 243/263 rewarded laser-off, 35/34 rewarded laser-on, 39/48 omission laser-off, 34/43 omission laser-on). Shaded regions show s.e.m. across trials. **d,** Violins quantify anticipatory licking (mean [-0.7,-0.2] s relative to reward; normalized as above) during all 4 conditions. GrC inhibition trials abolished anticipatory licking (laser off vs on: VR rewarded, VR omission, Reach rewarded all p<10^-6^; Reach omission p=0.00001; two-sided Mann-Whitney U test; **Methods**).

**Extended Data Figure 3.**
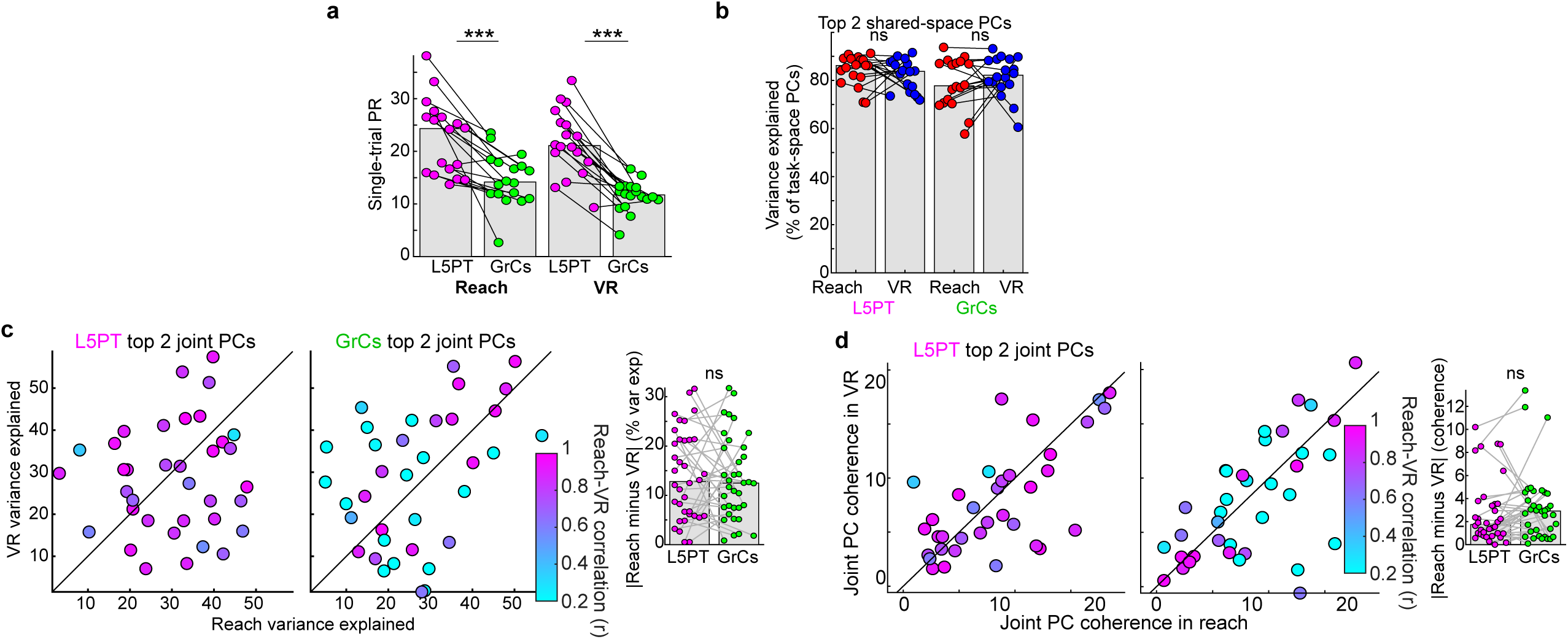
| Comparison of GrC and L5PT dynamics. **a,**Effective dimensionality (participation ratio [PR]) computed on single-trial data for each task. For each session, we concatenated all rewarded trials into a Time×Cells matrix and computed PCA and PRs, which were significantly lower for GrCs than for L5PT (both p=0.0003, Wilcoxon signed-rank test), thus as in Fig. 2l**,m** qualitatively inconsistent with a traditional high-rank GrC expansion (18 matched VR and Reach session pairs from 9 mice). **b-d,** Goodness of fit for joint-task PCA data shown in Fig. 3. **b,** Variance explained by the top 2 joint-task PCs as a percentage of the variance explained by the top 2 “task space” PCs. This metric controls for the possibility that the low cross-task correlations in GrCs (Fig. 3u) were simply an artifact of poor joint-subspace fitting. However, joint PCs captured the vast majority of variance captured by per-task PCA for both populations (L5PT ∼85%, GrCs ∼80%), and this ratio was indistinguishable between tasks for both cell types (L5PT: p=0.4; GrCs: p=0.1, Wilcoxon signed-rank test). **c,d,** Joint PC quality variation between tasks. Variance explained **(c)** and coherence **(d)** for the top 2 joint-task PCs from all sessions. To explicitly test if GrCs suffered from a harsher cross-task compromise than L5PT, we calculated the absolute difference between the Reach and VR metrics (distance from the unity line, insets). Both L5PT and GrCs showed comparable deviation from unity (insets; Variance: p = 0.6; Coherence: p = 0.1, Wilcoxon signed-rank test). By contrast, cross-task correlations (color scale) were profoundly lower for GrCs—demonstrating that GrC decorrelation cannot be attributed to lower joint PC quality or asymmetric subspace fitting. Coherence is quantified as the ratio of PC variance to the variance of a random projection (18 matched VR and Reach session pairs from 9 mice).

**Extended Data Figure 4.**
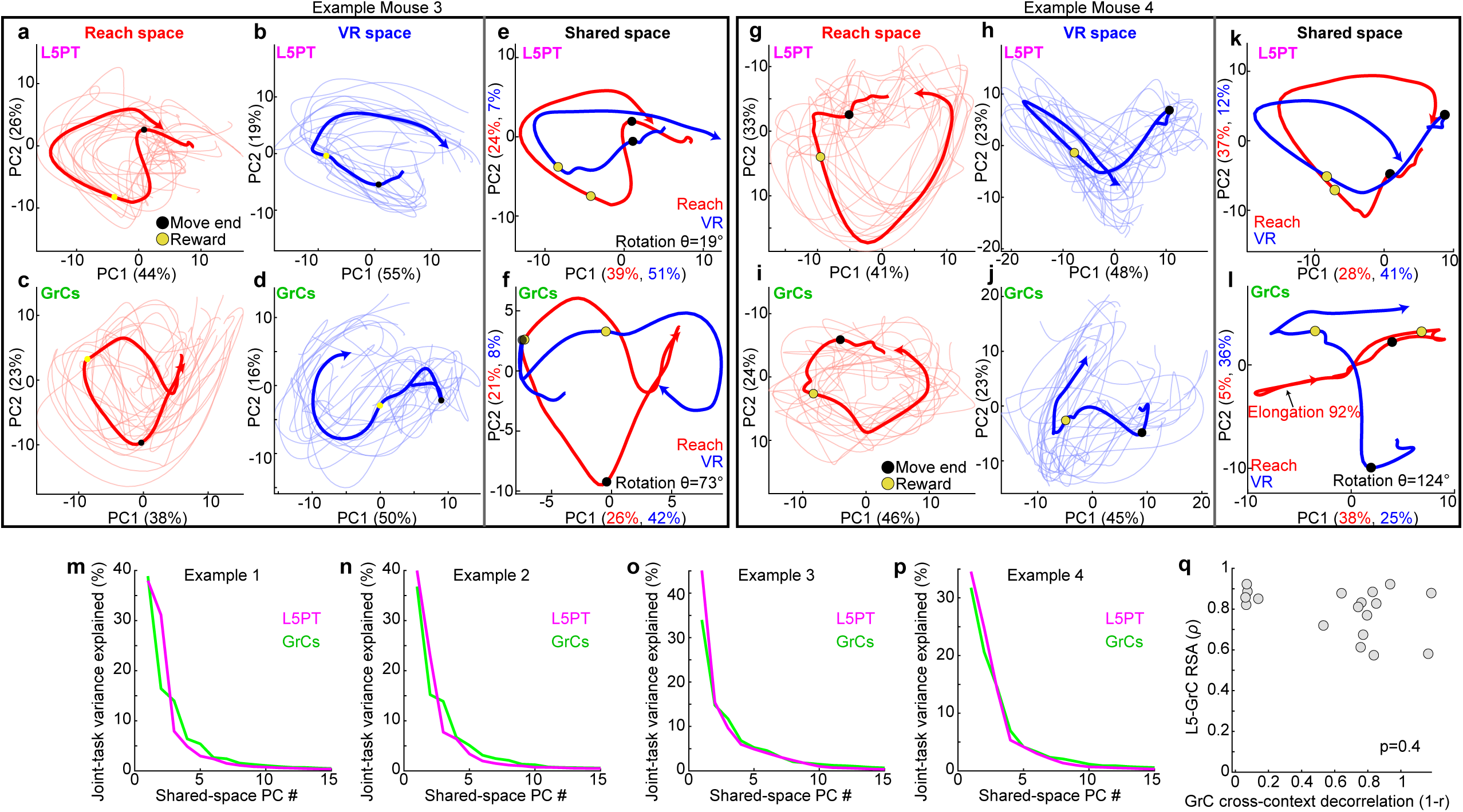
| Additional examples of GrC trajectory reorientations. **a–l,** Low-dimensional trajectory analysis for two additional mice (Mouse 3 and Mouse 4), distinct from those shown in Fig. 4. **a–f,** Example Mouse 3: A session characterized by strong GrC rotational separation. **a–d,** Independent task-space projections show smooth cyclic motifs in both cell types and tasks. **e,** In the joint-task shared space, L5PT trajectories remain aligned (trajectory reuse, rotation∼19**°**). **f,** GrC trajectories, however, undergo a large relative in-plane rotation (rotation∼73**°**), orthogonalizing the contexts while maintaining the cyclic structure. **g–l,** Example Mouse 4: A session exhibiting concurrent GrC in-plane rotation and out-of-plane separation. **g–j,** Independent projections. **k,** L5PT trajectories remain aligned in shared space (rotation∼17**°**). **l,** GrC trajectories show a complex transformation: the Reach trajectory (red) is both strongly rotated (rotation∼124**°**) relative to VR but also significantly elongated (∼92%), appearing effectively “collapsed” along PC2. This illustrates that context separation in GrC trajectories often manifests as both in-plane and out-of-plane reorientations. For visualization only, means and single trials were Gaussian smoothed with σ=0.1 and 0.2 s respectively. **m-p,** Scree plots of the joint-task variance explained by each shared-space PC for Examples 1-4. **q,** Cross-context GrC trajectory decorrelation versus L5-GrC similarity (dots denote session pairs, *ρ* values from Fig. 4q). Even when GrCs strongly decorrelated contexts, L5-GrC representational similarity usually remained high (Spearman’s *ρ=* –0.21, p=0.4, two-sided. 18 matched VR and Reach session pairs from 9 mice).

**Extended Data Figure 5.**
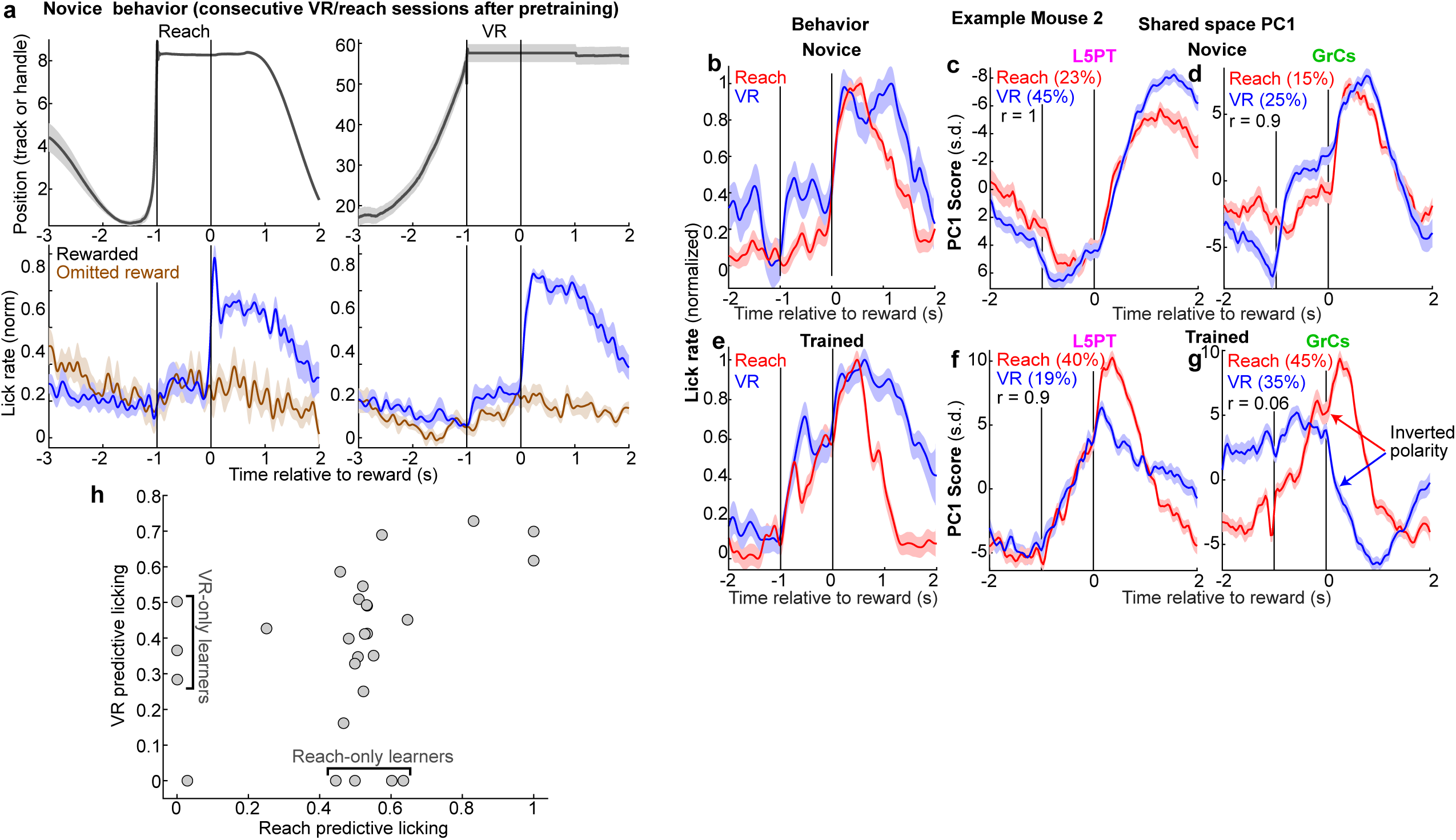
| Additional characterization of learning. **a,**Cohort-averaged Novice behavior (analogous to Fig. 1) in Reach (left column) and VR (Right column). “Novice” data are derived from consecutive VR/Reach training days, e.g., Reach on Day N and VR on Day N+1 acquired after pre-training. Top: Position trajectories (Reach robotic handle position or VR running sphere position). Bottom: Lick rates on rewarded and omitted reward trials, showing primarily reactive licking strategies. Traces represent the average of trial-averages from 9 paired novice Reach-VR sessions from 9 mice. Shaded regions show s.e.m. across sessions. **b-g**, A second representative example mouse showing learning-related changes in licking behavior and in both L5PT and GrC joint-task PC1 activity in both Reach and VR (analogous to Fig. 5a**-f**; Novice: 51 Reach/76 VR trials; Trained: 91 Reach/43 VR trials). Shaded regions show s.e.m. across trials. **h,** Comparison of predictive licking performance in each task independently (quantified per-task as in Fig. 5). Brackets highlight 3 sessions with predictive licking in VR but not in Reach, and conversely 4 sessions with significant predictive licking in Reach but not in VR, demonstrating that during intermediate learning stages, animals sometimes acquire predictive behavior in only one task, rather than consistently learning both tasks in lockstep (26 session pairs; 17 trained, 9 novice).

**Extended Data Figure 6.**
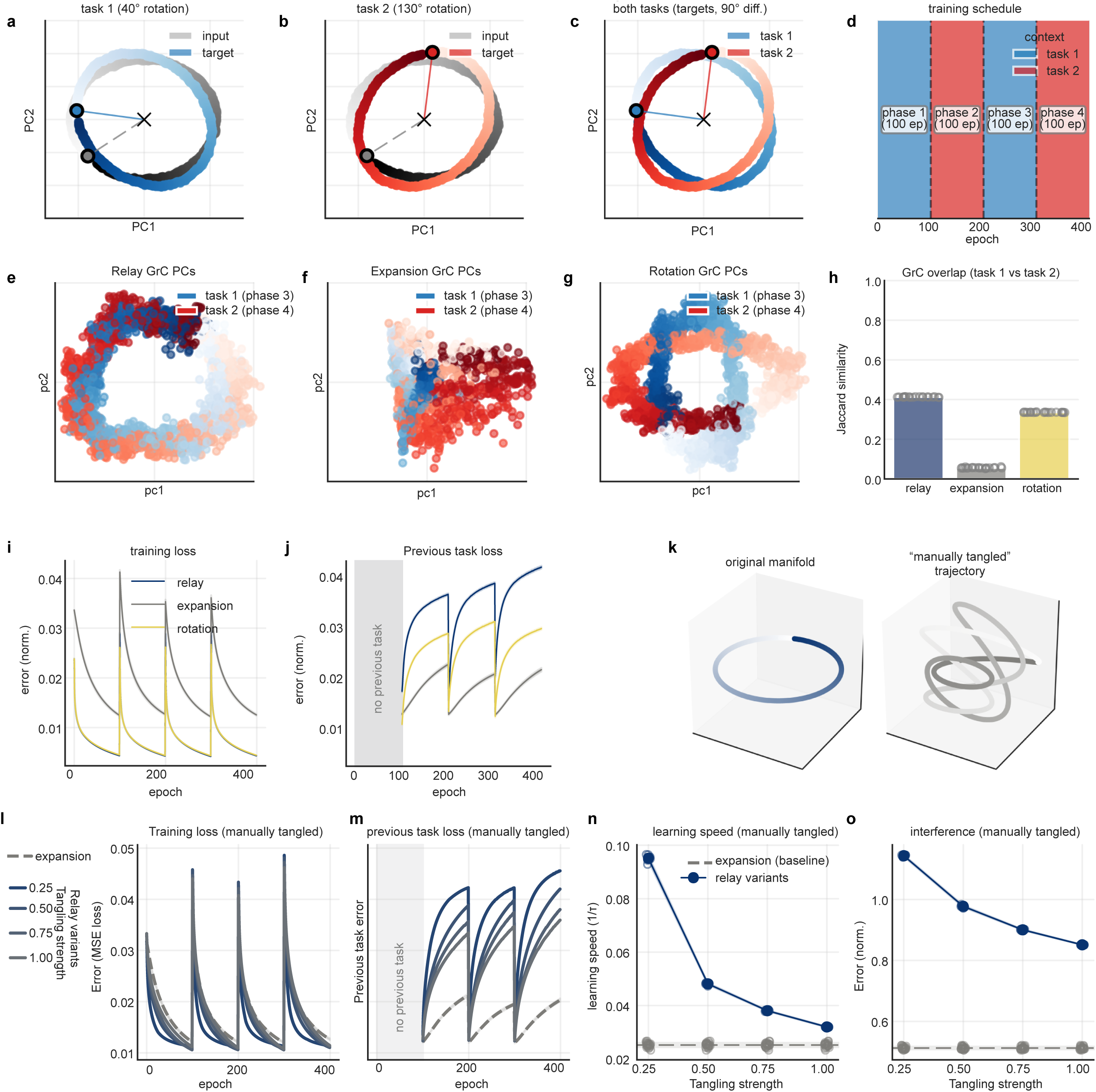
| Characterization of simulated architectures and the geometric trade-off between learning speed and interference. **a-c,** State-space geometry of the simulated dynamic prediction tasks, illustrating input manifolds (grey) and temporally shifted target predictions (colored) for Task 1 (40° rotation, **a**), Task 2 (130° rotation, **b**), and both tasks combined (**c**). Noiseless inputs are shown for visualization purposes; all simulations used noisy inputs. **d,** The alternating training schedule, comprising 100-epoch phases. **e-h,** Principal component projections of GrC representations during alternating tasks. **e**, The Cortical Relay model preserves overlapping geometry. **f**, The High-Rank Expansion model reveals a spectrally whitened high-dimensional geometry (axes scaled independently to each model to visualize structure despite sparsity-induced variance reduction). **g**, The Low-Rank Rotation model geometrically separates the trajectories while preserving smooth low-dimensional geometry. **h,** Active GrC population overlap measured by Jaccard similarity. Due to dense coding, both Relay and Rotation models recruit largely overlapping populations. In contrast, the Expansion model sparsifies the representation, resulting in near-zero overlap. This confirms the Rotation model mitigates interference through geometric alignment rather than physical population partitioning (dots show 20 random network initializations). **i,** Training loss over epochs for the three GrC architectures. Note the rapid convergence of the Relay and Rotation models relative to the Expansion model. **j,** Previous task loss (Task 2 loss when trained on Task 1, and vice versa) over epochs. The Relay model suffers from catastrophic interference immediately following task switches. The Rotation model maintains lower error on the inactive task, comparable to the Expansion model. **k-o,** Isolating the geometric effects of smoothness. **k**, To test whether the loss of smoothness—beyond the effects of sparsity—drives the learning-interference trade-off, structurally intact (left) or “manually tangled” (right) manifolds were fed into the dense Relay architecture. **l-o**, Increasing the geometric tangling strength within the Relay model progressively degraded training loss (**l**) and learning speed (**n**), while simultaneously reducing previous task loss (**m**, **o**). At maximum tangling, the dense Relay model’s performance morphed to match the sparse Expansion model baseline more closely (dashed grey lines). Dots in **h,n,o** depict 20 random network initializations.

**Extended Data Figure 7.**
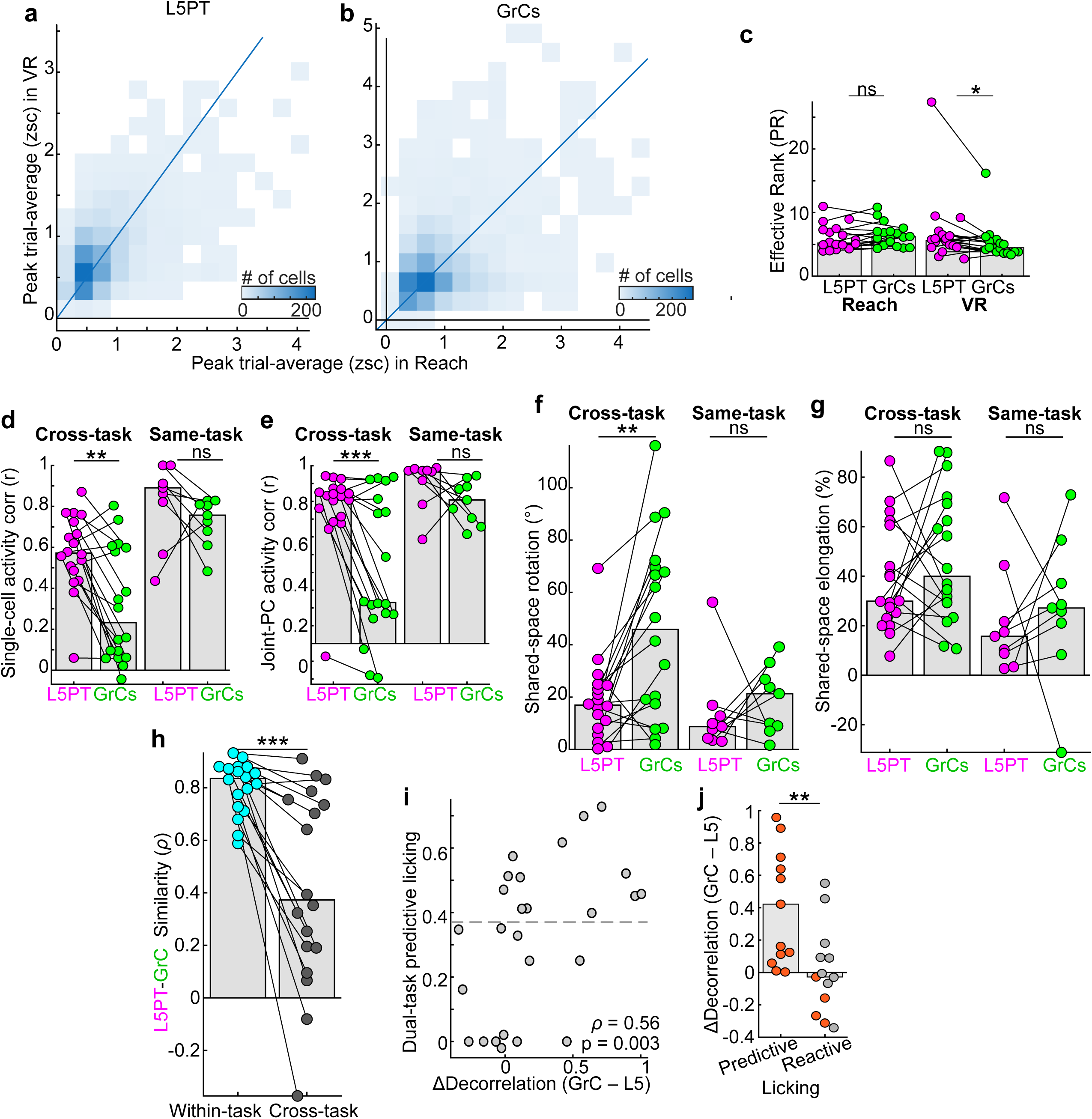
| Findings are robust to cell tracking and inclusion criteria. **a,b,** Joint histograms of activity level (peak trial-averaged z-scored fluorescence) across tasks for all cross-task-matched L5PT neurons (**a**) and GrCs (**b**). Thresholding these populations to identify nearly task-specific neurons (>0.7 zsc in one task but <0.4 zsc in the other) identified only 8.1% of L5PTs and 7.7% of GrCs as putatively task-specific by this metric. This was consistently, but only slightly, higher than in cross-day same-task negative control comparisons (4.8% of L5PTs and 5.1% of GrCs). Thus, neurons successfully matched across tasks were overwhelmingly likely to be comparably active in both, with no difference between cell types. **c-j**, Robustness of representational findings to cell selection criteria. To ensure our central conclusions were not driven by the exclusion of task-unreliable neurons, we replicated key analyses utilizing all tracked cells, regardless of their split-half reliability. The central representational results remained nearly identical to the filtered dataset. Specifically, in the unfiltered population comparing L5PT to GrCs: effective dimensionality (**c**) was 5.2 vs 6.2 in Reach (p=0.2) and 5.6 vs 4.5 in VR (p=0.03; compared to 4.1/5.0 in Reach and 4.3/4.1 in VR in Fig. 2m); single-cell correlations (**d**; cross-task: p=0.002; same-task: p=0.2) cross-task were r=0.57 vs r=0.23 (compared to 0.58/0.23 in Fig. 3o); shared-space PC1-2 correlations (**e;** cross-task: p=0.001; same-task: p=0.2) cross-task were r=0.83 vs 0.23 (compared to 0.83/0.24 in Fig. 3u); trajectory rotation (**f**; cross-task: p=0.002; same-task: p=0.4) cross-task was 17° vs 46° (compared to 19°/54° in Fig. 4m); elongation (**g**; cross-task: p=0.4; same-task: p=0.5) cross-task was 30% vs 40% (compared to 26%/46% in Fig. 4n). Within-task L5-GrC RSA similarity (**h**, p=0.0002) was *ρ*=0.83 (identical to Fig. 4q). Excess GrC decorrelation correlated with predictive licking (**i**) at *ρ*=0.56 (compared to 0.6 in Fig. 5i), and median decorrelation in predictive versus reactive sessions (**j**, p=0.003) was 0.42 vs -0.03 (compared to 0.44/-0.004 in Fig. 5j). With the exception of the elongation score— where the large population of task-unreliable cells elevated the isotropic variance floor, obscuring the geometric signal—the representational and geometric findings were robust to cell selection criteria. All quantifications across 18 cross-task (9 mice) and 9 same-task (7 mice) session pairs. All paired and unpaired comparisons used the Wilcoxon signed-rank or two-sided Mann-Whitney U test.

